# Iron-dependent reprogramming of damage-associated peptide receptor signaling coordinates immunity and phosphate stress adaptation

**DOI:** 10.64898/2026.03.09.710003

**Authors:** Natsuki Tsuchida, Tae-Hong Lee, Maxmilyand Leiwakabessy, Kota Yamashita, Kei Hiruma, Kentaro Okada, Taishi Hirase, Shigetaka Yasuda, Yuniar Devi Utami, Salvatore Cosentino, Hirotaka Ariga, Miki Fujita, Min Fey Chek, Taishi Umezawa, Yusuke Saijo

**Affiliations:** Graduate School of Science and Technology, Nara Institute of Science and Technology, 630-0192, Ikoma, Japan; Graduate School of Bio-Applications and Systems Engineering, Tokyo University of Agriculture and Technology, 184-8588, Tokyo, Japan; Informatics Platform, Center for Cancer Immunotherapy and Immunobiology, Kyoto University Graduate School of Medicine, 606-8501, Kyoto, Japan; Research Center of Genetic Resources, National Agriculture and Food Research Organization, 305-8602, Ibaraki, Japan; Mass Spectrometry and Microscopy Unit, RIKEN Center for Sustainable Resource Science, 305-0074, Ibaraki, Japan; Department of Life Sciences, Graduate School of Arts and Sciences, The University of Tokyo, 153-8902, Tokyo, Japan

## Abstract

Plant adaptation to inorganic phosphate (Pi) limitation entails extensive developmental and metabolic reprogramming, collectively termed the phosphate starvation response (PSR). How plants balance nutrient acquisition with immune competence remains unresolved. Here, we show that PSR selectively remodels pattern-recognition receptor (PRR) signaling in *Arabidopsis thaliana*. Membrane proteomics reveals increased abundance of the Pep receptor PEPR1 under Pi deficiency, whereas most receptor kinases, including the microbial PRRs FLS2 and CERK1, are reduced. Immunoblotting further shows increased PEPR2 abundance. Consistently, Pep-triggered responses are strongly potentiated under Pi depletion, while flagellin- and chitin-induced responses are not, indicating selective PEPR sensitization rather than global immune suppression. This sensitization requires the PSR-associated ferroxidases LPR1/LPR2, NADPH oxidases RBOHD/RBOHF, and transcription factor WRKY33, linking iron-dependent redox remodeling to immune amplification. Beyond inducible defense, PEPRs promote basal PSR transcriptional reprogramming and growth under Pi limitation. Loss of *PEPR1/PEPR2* compromises bacterial resistance and alters root-associated microbiota assembly under nutrient deficiency. Together, PEPRs exhibit context-dependent dual functionality, promoting either immunity or nutrient stress adaptation, potentially in response to distinct PROPEP-derived ligands, thereby supporting nutrient acquisition without compromising immune vigilance. These findings provide a framework for how nutritional status selectively remodels PRR signaling to coordinate immunity with nutrient stress adaptation.

## Introduction

Plants dynamically adjust the immune system to maintain nutrient homeostasis and accommodate beneficial microbial associations. Activation of defense responses often constrains growth and nutrient acquisition, reflecting a fundamental growth–defense trade-off^1–5^. This trade-off becomes particularly acute under nutrient limitation, where plants must optimize resource allocation for nutrient acquisition while preserving protection against pathogens. How immune competence is sustained without compromising nutrient stress adaptation remains a central unresolved question in plant biology.

Phosphorus (P) is an essential macronutrient that often limits plant productivity, as inorganic phosphate (Pi) readily forms insoluble complexes with metals such as iron (Fe) in soils. In response to Pi scarcity, plants activate the phosphate starvation response (PSR), largely governed by the MYB transcription factors PHOSPHATE RESPONSE1 (PHR1) and PHR1-LIKE1 (PHL1) ^6–8^. PSR encompasses transcriptional induction of high-affinity PHOSPHATE TRANSPORTER1 (PHT1) transporters, remodeling of root system architecture, and secretion of organic acids and phosphatases to mobilize Pi from the soil^9,10^. In parallel, primary root growth is inhibited through *PHR1*-independent, Fe-dependent redox processes in the root meristem. The ferroxidases LOW PHOSPHATE ROOT1 and 2 (LPR1/LPR2) oxidize Fe²□ to Fe³□, promoting Fe³□ accumulation, reactive oxygen species (ROS) production, and callose deposition. These processes restrict cell-to-cell movement of the transcription factor SHORT-ROOT, thereby impairing radial patterning and stem cell maintenance^11–13^. PSR also entails extensive rhizosphere remodeling to enhance P mobilization. Carbohydrate secretion and proton extrusion lead to rhizosphere acidification, which increases the solubility and availability of both Pi and Fe at the root-soil interface^14^. Such acidification suppresses immune signaling and bacterial resistance in the root^15^. Given that Fe availability and ROS are key modulators of immune signaling^16,17^, these observations suggest that PSR-driven changes in rhizosphere chemistry may reshape immune signaling capacity.

Beyond intrinsic adaptive responses, plants engage beneficial microbes to improve nutrient acquisition. Although *Arabidopsis thaliana* (hereafter Arabidopsis) does not establish arbuscular mycorrhizal symbioses, it associates with beneficial endophytic fungi such as *Colletotrichum tofieldiae*, which promotes Pi uptake under P limitation^18^. During beneficial interactions with *C. tofieldiae*, defense-related gene expression is attenuated, whereas a pathogenic relative triggers defense activation even under Pi deficiency^19^. Thus, immune outputs are dynamically modulated according to nutritional status and microbial lifestyle. However, the molecular mechanisms that preserve pathogen resistance under nutrient stress remain unclear.

Pattern-triggered immunity (PTI) in plants is initiated by cell-surface pattern-recognition receptors (PRRs) that detect microbe-or damage-associated molecular patterns (MAMPs/DAMPs)^20–22^. Canonical MAMP receptors in Arabidopsis include the leucine-rich repeat (LRR) receptor kinase (RK) FLAGELLIN-SENSITIVE2 (FLS2), which detects bacterial flagellin^23^, and the lysin-motif (LysM) RK CHITIN ELICITOR RECEPTOR KINASE1 (CERK1), which mediates perception of fungal chitin and bacterial peptidoglycan^24,25^. Activation of these receptors triggers conserved signaling cascades, involving ROS burst, mitogen-activated protein kinase (MAPK) activation, and extensive transcriptional reprogramming. Endogenous danger signals also elicit comparable PTI outputs through phytocytokine receptors^26,27^. The LRR-RKs PEP1 RECEPTOR1 (PEPR1) and PEPR2 perceive plant elicitor peptides (Peps) derived from PROPEP precursors, while RECEPTOR-LIKE KINASE7 (RLK7) recognizes PAMP-induced peptides (PIPs) derived from PREPIP precursors^28–31^. These receptors amplify and propagate defense signaling downstream of primary MAMP perception and effector-triggered immunity under nutrient sufficiency ^32–35^. PEPR signaling can compensate for impaired immunity when depletion of the shared coreceptor BRI1-ASSOCIATED RECEPTOR KINASE1 (BAK1) compromises BAK1-dependent PRR pathways^36^. In addition to immune amplification, PEPR2 has been implicated in root meristem differentiation under low Pi via CLAVATA3/EMBRYO SURROUNDING REGION-RELATED 14 peptide signaling in conjunction with the LRR-RK CLAVATA2^13^, suggesting that PEPR signaling can intersect with nutrient-responsive developmental pathways.

Recent studies established that PSR antagonizes PTI, showing that FLS2-mediated signaling is attenuated under Pi starvation in a *PHR1*-or *PHT1*-dependent manner^37–39^. These findings have been interpreted as evidence for a global nutrient–immunity trade-off in which Pi starvation suppresses immune signaling. However, whether Pi limitation uniformly dampens PRR pathways or instead selectively reconfigures immune receptor abundance and signaling capacity has remained unresolved. Here, we demonstrate that Pi deficiency does not globally suppress PRR signaling but differentially modulates specific immune pathways. Quantitative membrane proteomics reveals preferential retention or increased abundance of a subset of RKs, including PEPR1 and RLK7, whereas several MAMP receptors are reduced. Moreover, only specific downstream defense outputs of retained receptors are enhanced, indicating pathway- and output-selective potentiation rather than uniform amplification of PRR signaling. Genetic and transcriptomic analyses further indicate that PEPR signaling contributes not only to inducible defense but also to basal PSR-associated transcriptional reprogramming and plant growth. Together, our findings reveal that Pi limitation selectively remodels specific PRR pathways and enhances Pep–PEPR responsiveness, providing a mechanistic framework for coordinating nutrient adaptation with immune competence.

## Results

### Pi deficiency differentially alters the abundance of plasma membrane receptors

To determine whether Pi starvation affects the abundance of PRRs, we performed quantitative proteomic analysis of endogenous plasma membrane proteins from Arabidopsis seedlings grown under Pi-sufficient and Pi-deficient conditions (Fig. 1a). Across conditions, 116 plasma membrane–localized receptor proteins were identified, spanning multiple receptor classes, including LRR-RKs, cysteine-rich repeat receptor kinases (CRKs), LysM-RKs, lectin RKs, CrRLK1L family members, and receptor-like proteins (RLPs) (Fig. 1b; Supplementary Fig. 1; Supplementary Data 1). Among these were previously described PRRs such as PEPR1, RLK7, CERK1, and FLS2, whereas PEPR2 was not detected above the analytical threshold.

**Fig. 1:**
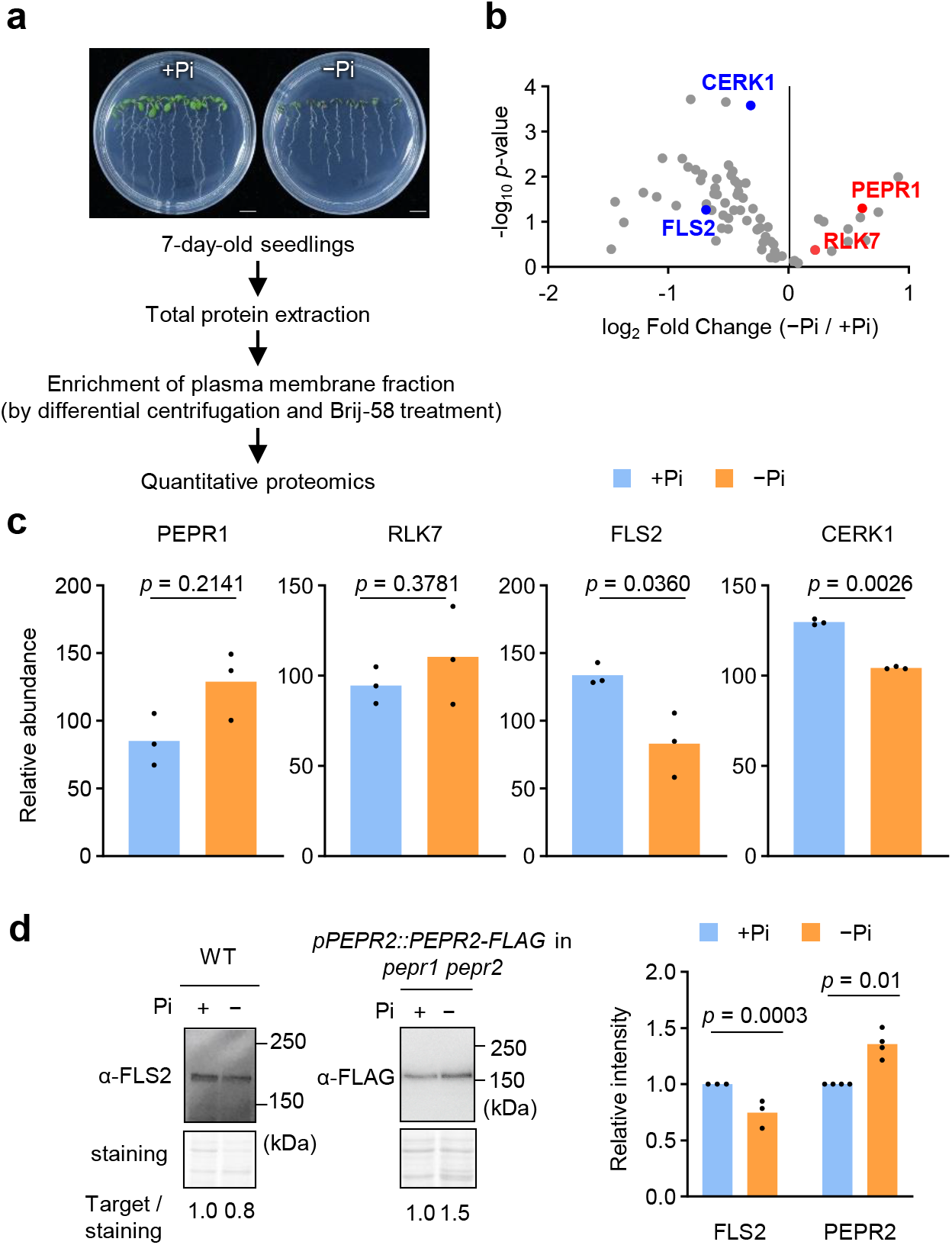
Pi deficiency differentially alters abundance of plasma membrane receptors. **a**, Workflow for proteomic analysis of plasma membrane-enriched fractions from 7-day-old seedlings grown under Pi-sufficient (+Pi, 625 µM) or Pi-deficient (−Pi, 10 µM) conditions. **b**, Volcano plot showing changes in receptor protein abundance in wild-type (WT) seedlings. The x-axis indicates the log□average abundance ratio, and the y-axis indicates the negative log□□-transformed, adjusted *p*-values. **c**, Relative abundance of the PRR proteins under +Pi and −Pi conditions. Values are scaled protein abundance calculated using Proteome Discoverer. Data are means, *n* = 3 biological replicates. *p-*values were calculated using two-sided Student’s *t*-test. **d**, Immunoblot analysis of endogenous FLS2 in WT seedlings and PEPR2-FLAG in *pPEPR2::PEPR2-FLAG / pepr1 pepr2* seedlings grown under +Pi and −Pi conditions. SYPRO Ruby staining served as a loading control (bottom). Band intensities were quantified using ImageJ and normalized to the corresponding +Pi controls, which were set to 1. Relative band intensities are shown below the immunoblots. Data are means of biological replicates, *n* = 3 for FLS2 and *n* = 4 for PEPR2–FLAG. Statistical significance between +Pi and −Pi conditions was determined using two-way ANOVA followed by Sidak’s multiple-comparisons test.

Comparative quantification revealed a non-uniform response to Pi deprivation. The abundance of PEPR1 and RLK7 was maintained or increased under Pi-deficient conditions, whereas numerous other receptors, including CERK1 and FLS2, were reduced relative to Pi sufficiency (Fig. 1c). Independent immunoblot analysis of FLS2 confirmed its reduced abundance under Pi deficiency, validating the proteomic results (Fig. 1d). Because PEPR1 and PEPR2 jointly mediate Pep perception, we next assessed whether PEPR2, which was not detected in the proteomic analysis, was also regulated by Pi availability. Immunoblot analyses of plasma membrane protein fractions from transgenic lines expressing *PEPR2-FLAG* under its native promoter revealed increased PEPR2 abundance under Pi-deficient conditions (Fig. 1d). Together, these results show that Pi deficiency differentially alters the abundance of specific membrane-resident immune receptors, increasing or maintaining both PEPRs and RLK7 while reducing several other PRRs.

Interestingly, transcript levels of *PEPR1*, *PEPR2*, *FLS2*, and *CERK1* were all elevated under low Pi conditions (Supplementary Fig. 2a), despite their divergent protein abundance profiles. Thus, selective receptor retention or reduction under Pi deficiency cannot be explained solely at the mRNA level, indicating substantial post-transcriptional regulation or differential protein stability. Among these genes, only *PEPR1* harbors a canonical PHR1-binding site^6^ in its regulatory DNA sequences (Supplementary Fig. 2b). Consistently, *PEPR1* expression was induced under low Pi partly in a *PHR1/PHL1*-dependent manner (Supplementary Fig. 2c), indicating direct integration of *PEPR1* transcription into the core PSR regulatory network.

### Pi deficiency potentiates PEPR signaling through selective amplification of downstream outputs

To determine how Pi availability influences PRR signaling competence, we compared immune responses mediated through PEPR1/PEPR2 and CERK1 pathways under Pi deficiency. Given that FLS2 signaling is attenuated under low Pi^39^, comparison with CERK1 allowed us to assess whether Pi limitation broadly suppresses PRR pathways across RK classes or differentially modulates specific PRR pathways.

Root transcriptomes were analyzed following Pep1 or chitin treatment under Pi-sufficient and Pi-deficient conditions (Supplementary Data 2 and 3). Pi deficiency extensively reprogrammed the Pep1-responsive transcriptome, preferentially enhancing the induction of a subset of Pep1-responsive genes while also modifying Pep1-mediated gene repression (Supplementary Data 2). Among the top 150 genes induced by each elicitor under Pi sufficiency, many Pep1-induced genes exhibited further enhancement under Pi deficiency (Fig. 2a). By contrast, most chitin-induced genes showed attenuated induction, with only a minority retaining comparable induction (Fig. 2b). Thus, Pi limitation preferentially amplifies specific PEPR-dependent transcriptional outputs rather than PRR-triggered transcription generally. Pep1-responsive genes were further categorized according to whether their induction or repression was enhanced or attenuated under Pi deficiency (Supplementary Data 4).

**Fig. 2:**
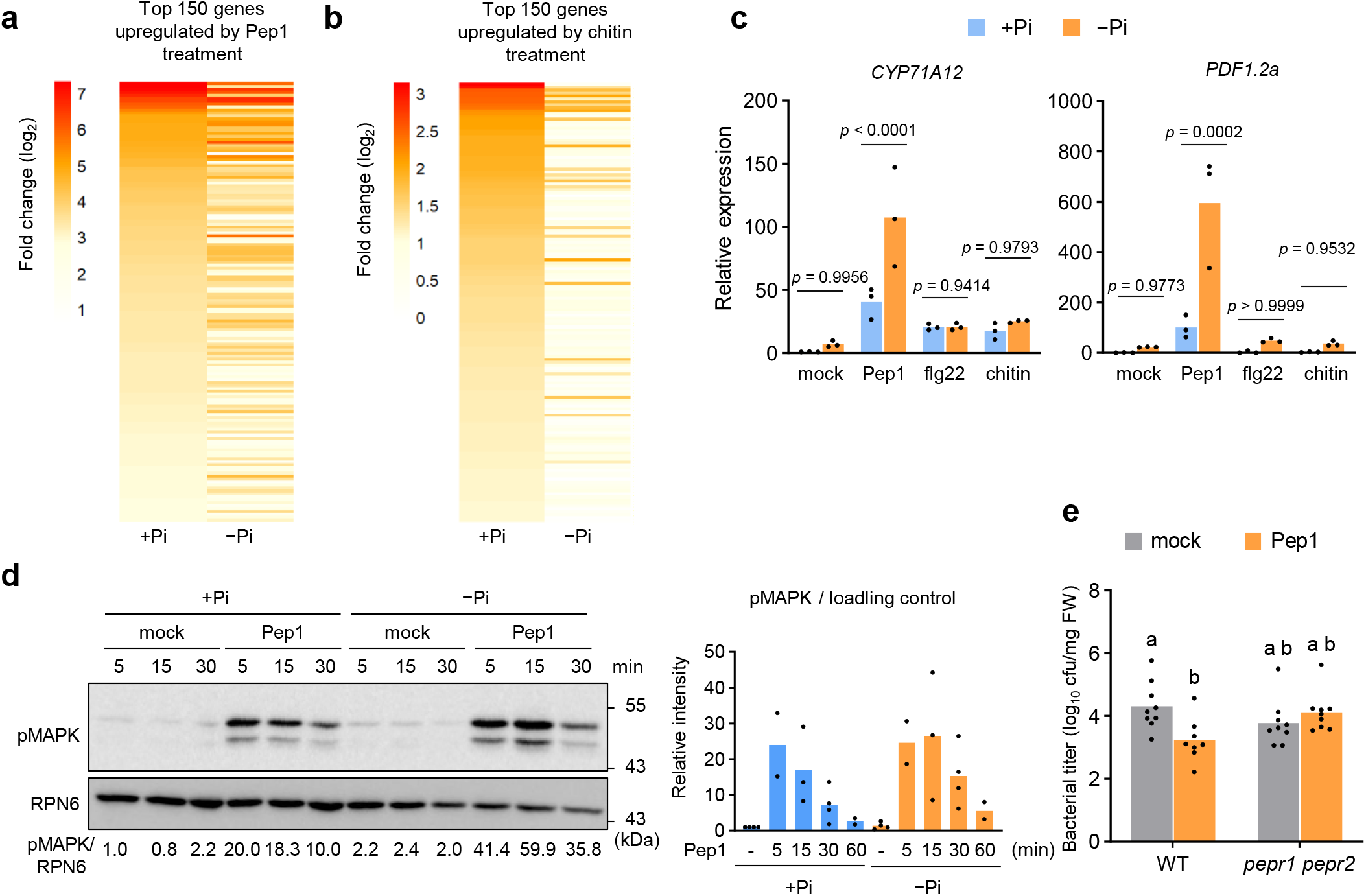
Pi deficiency potentiates PEPR signaling through selective amplification of downstream outputs. **a,b,** Fold changes of the top 150 genes induced by Pep1 (a) or chitin (b) treatment for 6 h under Pi-sufficient conditions (+Pi, 625 µM). Values represent log_2_ fold change relative to the mock-treated controls under Pi-sufficient conditions (+Pi, 625 µM) or Pi-deficient conditions (−Pi, 10 µM). **c**, *CYP71A12* and *PDF1.2a* transcript levels in WT roots grown under +Pi or −Pi conditions and treated with water (mock), 1 µM Pep1, 1 µM flg22, or 1 mg ml□¹ chitin for 6 h. Data are means, *n* = 3 biological replicates. Statistical significance between +Pi and −Pi conditions was determined using two-way ANOVA followed by Sidak’s multiple-comparisons test. **d**, Immunoblot analysis of Pep1-induced MAPK phosphorylation in WT roots grown under +Pi or −Pi conditions and treated with water or 1 µM Pep1 for the indicated times. RPN6 served as a loading control (bottom). Representative blots are shown. Numbers below the blots indicate phosphorylated MAPK band intensities normalized to RPN6 and expressed relative to the +Pi mock sample at 5 min (set to 1). Data are means, with *n* = 4, 2, 3, 4, and 2 biological replicates at 0, 5, 15, 30, and 60 min, respectively. **e**, Growth of *Pto* DC3000 in WT and *pepr1 pepr2* plants grown in nutrient-deficient soil after mock or Pep1 treatment. Data were pooled from four independent experiments and are presented as means, with *n* = 8 and *n* = 9 biological replicates for WT Pep1 and the other groups, respectively. Different letters indicate significant differences (*p* < 0.05; two-way ANOVA followed by Tukey’s multiple-comparisons test).

Quantitative RT–PCR (qRT-PCR) of the defense markers *CYP71A12* and *PDF1.2a*^40,41^ confirmed that Pep1-triggered gene expression was significantly enhanced in Pi-depleted roots, whereas responses to flg22 or chitin were unchanged or attenuated (Fig. 2c), consistent with reduced FLS2 signaling under low Pi^39^. Pep3-triggered responses, mediated specifically by PEPR1^29^, were similarly enhanced (Supplementary Fig. 3a). Consistent with the maintenance of RLK7 abundance, PIP2-induced *CYP71A12* expression was also enhanced under Pi deficiency (Supplementary Fig. 3b). This response was reduced, but not abolished, in *rlk7*, suggesting that RLK7 contributes to PIP2 signaling but may not be its sole receptor under these conditions. By contrast, SCOOP12-induced immune responses were not enhanced (Supplementary Fig. 3c), demonstrating that Pi deficiency does not generally potentiate signaling elicited by endogenous peptides but instead acts on specific PRR pathways.

Pep1-induced expression of *CYP71A12,* but not *PDF1.2a*, was further elevated in plants constitutively expressing PEPR1 under the Cauliflower mosaic virus 35S promoter (Supplementary Fig. 3d). These results indicate that increased receptor abundance contributes to the amplification of selected Pep-responsive outputs but is insufficient to account for the full potentiation of downstream signaling under Pi deficiency.

Time-course analyses revealed sustained amplification of Pep1-induced marker gene expression from 2 h to 10 h after treatment (Supplementary Fig. 3e), accompanied by stronger MAPK activation (Fig. 2d) and increased extracellular accumulation of PROPEP3-derived peptides (Supplementary Fig. 3f) under Pi deficiency. Enhanced Pep1 responsiveness persisted in *bak1-4* plants (Supplementary Fig. 3g), and BAK1 protein levels were unchanged between the two Pi conditions (Supplementary Fig. 1, 3h). Pi-dependent PEPR signaling potentiation is therefore distinct from the reinforcement of PEPR signaling associated with reduced BAK1 availability^36^. Moreover, Pep1-induced PEPR1–BAK1 complex formation and ROS burst were unaffected by Pi status (Supplementary Fig. 3h, i). Thus, Pi deficiency does not uniformly enhance PEPR signaling from receptor activation onward but selectively amplifies downstream signaling outputs, including MAPK activation and transcriptional reprogramming.

Pep1-induced immune responses were also enhanced in shoots under nutrient-deficient conditions (Supplementary Fig. 3j), indicating that PEPR sensitization is not restricted to roots. The sucrose-containing media used for seedling assays hampered bacterial inoculation; we therefore assessed disease resistance in mature leaves of soil-grown plants. Pep1 pretreatment significantly reduced *Pseudomonas syringae* pv. *tomato* (*Pto*) DC3000 growth in WT plants in all four experiments under nutrient deficiency, compared with two of four experiments under nutrient sufficiency (Fig. 2e; Supplementary Fig. 4a, b). Pep1-induced resistance under nutrient deficiency depended on PEPR1/PEPR2 (Fig. 2e; Supplementary Fig. 4a). By comparison, flg22 pretreatment robustly induced resistance under both nutrient conditions in both experiments tested (Supplementary Fig. 4b). These soil treatments differed in overall nutrient supply and did not isolate the contribution of Pi availability to disease resistance.

Collectively, these data demonstrate that Pi limitation differentially modulates the tested PRR pathways: it potentiates PEPR- and RLK7-mediated responses but does not enhance MIK2, FLS2, or CERK1 signaling. Within the PEPR pathway, potentiation is restricted to specific signaling branches rather than uniformly extending from receptor-proximal events to all downstream outputs. This pathway- and output-selective potentiation prompted us to investigate how distinct Pi-response modules reinforce PEPR-mediated immune signaling under nutrient stress.

### External Pi signaling via LPR1/LPR2 promotes PEPR-dependent immune potentiation under Pi deficiency

To dissect the regulatory basis of selective PEPR sensitization, we examined the contributions of local LPR1/LPR2-dependent iron signaling and systemic PHR1/PHL1-mediated PSR^6,8,11,42^. We compared root transcriptomic responses to Pep1 in WT, *phr1 phl1*, *lpr1 lpr2*, and *pepr1 pepr2* seedlings after 2 h and 10 h treatment under Pi-sufficient and Pi-deficient conditions (Supplementary Data 5 and 6). Differential expression analysis (log□ fold change > 1, FDR < 0.05) identified 1,511 and 1,288 PEPR-dependent Pep1-inducible genes under Pi-sufficient and Pi-deficient conditions, respectively, defined as transcripts significantly upregulated in Pep1-treated WT compared to Pep1-treated *pepr1 pepr2* seedlings (Fig. 3a). Although Pi deficiency did not increase the number of these genes, the magnitude of their Pep1-induced responses was largely greater under Pi deficiency (Fig. 3a, b). This indicates quantitative amplification of PEPR-dependent transcription rather than expansion of the PEPR-regulated gene set.

**Fig. 3:**
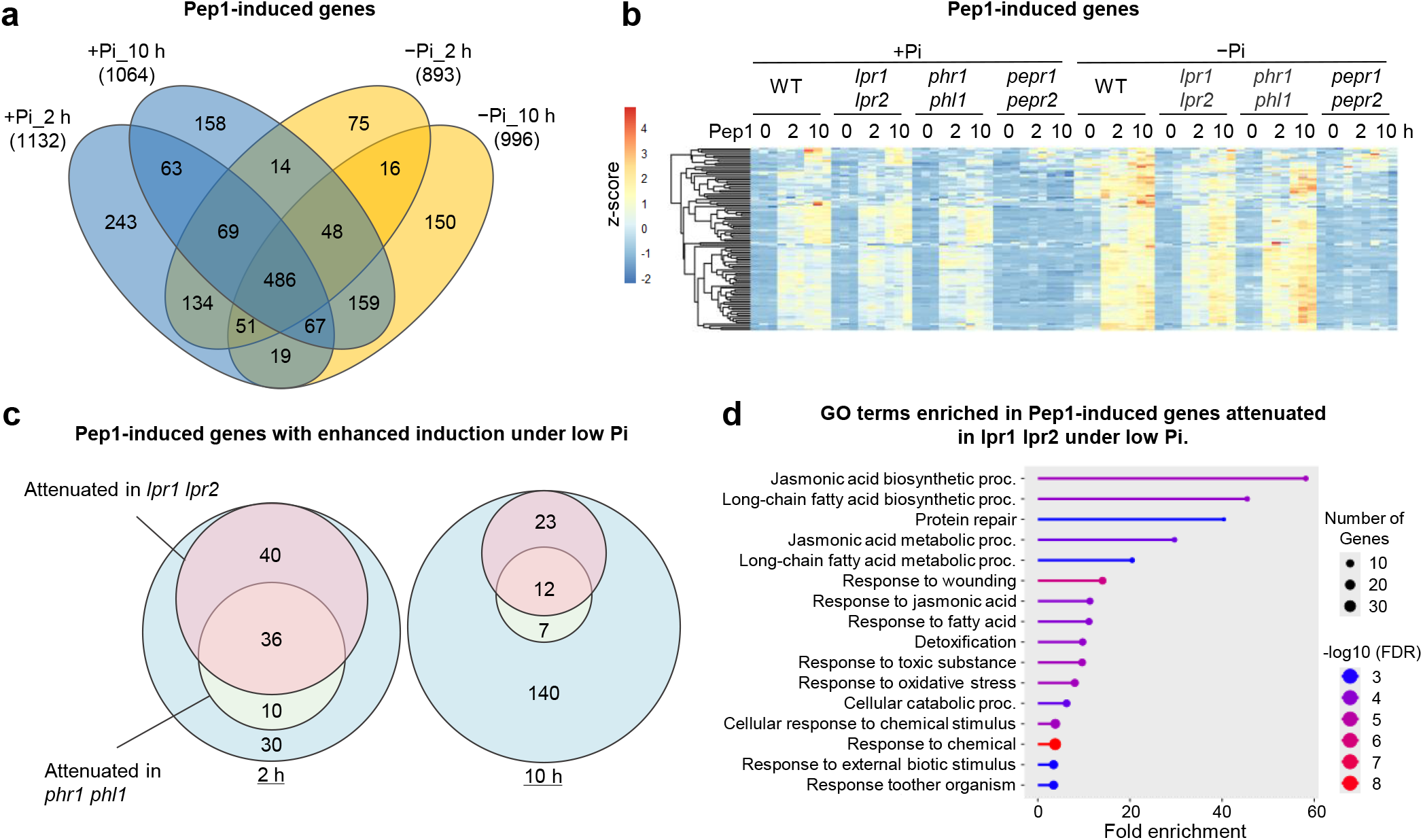
External Pi sensing via LPR1/LPR2 drives PEPR-dependent immune potentiation under Pi deficiency. **a**, Venn diagram showing Pep1-induced genes identified in WT seedlings under Pi-sufficient (+Pi) and Pi-deficient (−Pi) conditions at 2 h and 10 h after Pep1 treatment. Genes were defined as transcripts that were significantly upregulated in WT relative to the corresponding 0 h control and showed significantly higher expression in WT than in *pepr1 pepr2* at the same time point (log□FC > 1, FDR < 0.05 for both comparisons). **b**, Transcript profiles of PEPR-dependent Pep1-induced genes upregulated in WT roots at one or more time points (log□FC > 1, FDR < 0.05) and expressed at higher levels in WT than in *pepr1 pepr2* (log□FC > 1, FDR < 0.05). Genes assigned to the GO terms “defense response”, “response to biotic stimulus”, “response to wounding”, “response to stimulus”, and “response to stress” were selected for visualization. The heatmap shows z-scores calculated from log□(TPM + 1) values. **c**, Venn diagram showing Pep1-induced genes that were upregulated under low Pi relative to normal Pi conditions (log□FC > 0.5, FDR < 0.05) and downregulated in *lpr1 lpr2* or *phr1 phl1* relative to WT after Pep1 treatment. **d**, Representative GO terms enriched among genes downregulated in *lpr1 lpr2* relative to WT at 2 h after Pep1 treatment under low Pi conditions.

At 2 h, the enhanced Pep1 responses under Pi deficiency were compromised in both *lpr1 lpr2* and *phr1 phl1* (Fig. 3c, Supplementary Fig. 5). By 10 h, however, this impairment persisted predominantly in *lpr1 lpr2*, whereas the Pep1-inducible transcriptional profile in *phr1 phl1* largely resembled that of WT, with many genes exhibiting comparable or even stronger induction. Nevertheless, a subset of Pep1-responsive genes remained dependent on PHR1/PHL1 at 10h (Fig. 3c, Supplementary Fig. 5).

To identify genes whose Pep1 responsiveness was potentiated by Pi deficiency, we selected those showing greater Pep1 responsiveness under Pi-deficient than Pi-sufficient conditions (Δlog□FC > 0.5, FDR < 0.05) and assessed their induction in the two mutants. Loss of *LPR1/LPR2* impaired the potentiation of a substantially broader set of genes than loss of *PHR1/PHL1*. The genes affected in *lpr1 lpr2* were enriched for jasmonic acid (JA) biosynthesis and signaling and for defense responses (Fig. 3d), indicating that LPR1/LPR2-dependent iron signaling is a major contributor to the potentiation of JA-related transcriptional outputs in PEPR signaling under Pi deficiency. By contrast, the overall contribution of PHR1/PHL1 became more limited by 10 h, although PHR1/PHL1 remained required for the Pep1-induced expression of a subset of genes. Together, these data indicate that LPR1/LPR2-dependent iron signaling provides the predominant sustained input into PEPR-dependent immune potentiation under Pi deficiency, whereas PHR1/PHL1 exert a less persistent, counterbalancing regulatory influence.

### LPR1/LPR2-dependent iron-ROS signaling potentiates PEPR-mediated immunity under Pi deficiency

Because iron availability modulates LPR1/LPR2-dependent root responses to low Pi^43^, we examined whether iron levels influence PEPR-mediated defense activation under Pi deficiency. Under low Pi, Pep1-induced defense gene expression and MAPK activation were further enhanced by excess iron, whereas both responses were attenuated under iron-deficient conditions (Fig. 4a, b). In contrast, flg22- and chitin-triggered responses were not similarly potentiated by excess iron (Supplementary Fig. 6a, b), indicating that iron-dependent sensitization preferentially reinforces PEPR signaling. Despite this signaling amplification, Pep1-induced primary root growth inhibition occurred irrespective of iron or Pi status (Supplementary Fig. 6c), suggesting that PEPR-mediated defense activation and growth arrest are separable outputs.

**Fig. 4:**
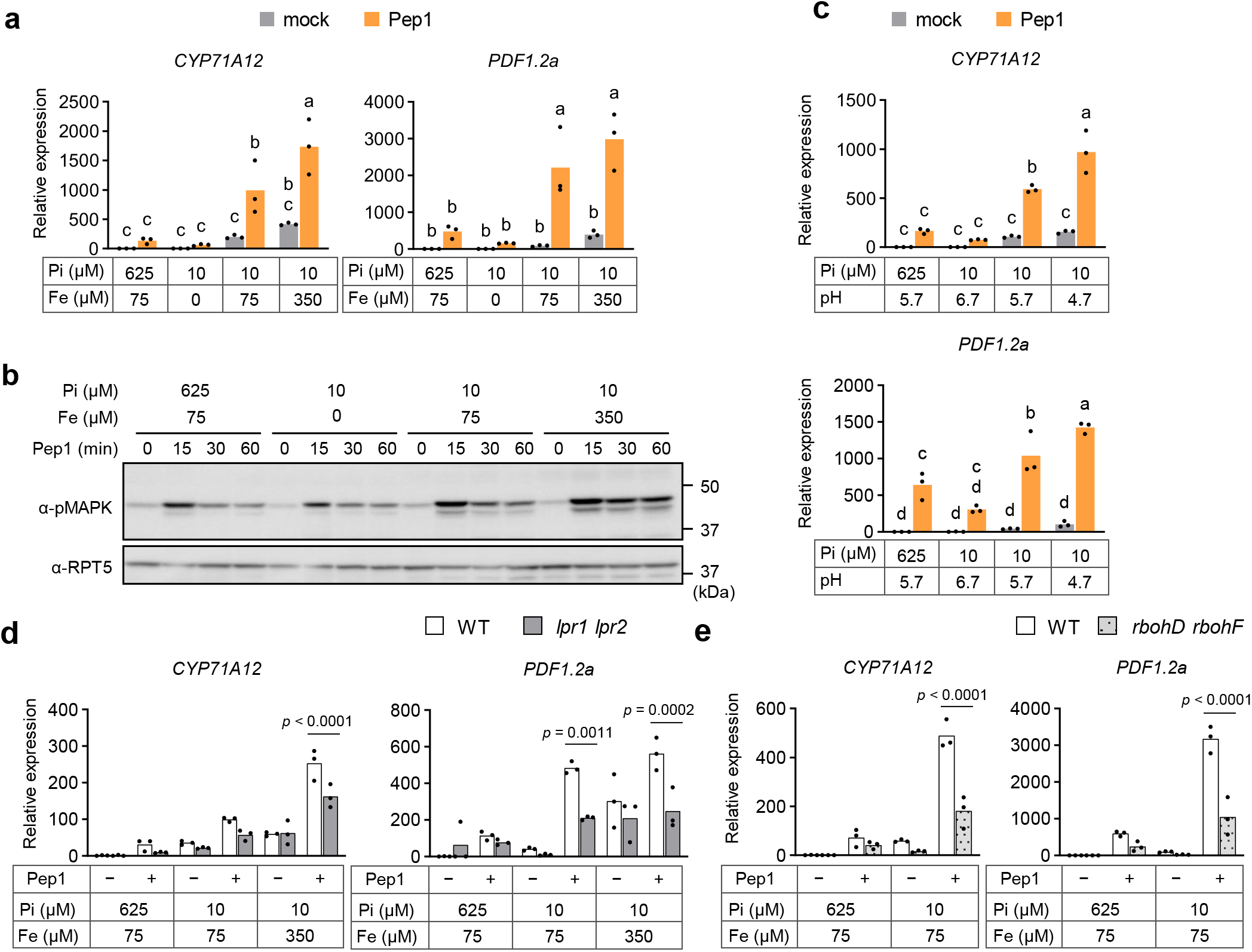
LPR1/LPR2-dependent iron-ROS signaling potentiates PEPR-mediated immune responses under Pi deficiency. **a**, *CYP71A12* and *PDF1.2a* transcript levels in WT roots grown under the indicated Pi and Fe conditions and treated with water (mock) or 1 µM Pep1 for 6 h. **b**, Immunoblot analysis of MAPK phosphorylation in WT roots grown under the indicated Pi and Fe concentrations and treated with 1 µM Pep1 for the indicated times. RPT5 is shown as a loading control (bottom). **c**, *CYP71A12* and *PDF1.2a* transcript levels in WT roots grown under the indicated Pi and pH conditions and treated with 1 µM Pep1 for 6 h. **d,e**, *CYP71A12* and *PDF1.2a* transcript levels in WT and *lpr1 lpr2* (**d**) or *rbohD rbohF* (**e**) roots grown under the indicated Pi and Fe conditions and treated with 1 µM Pep1 for 6 h. For transcript analyses (**a,c–e**), data are means, *n* = 3 biological replicates. In **a,c**, different letters indicate significant differences (*p* < 0.05; two-way ANOVA with Tukey’s multiple comparisons test). In **d,e**, *p* values for significant comparisons are indicated in the figure. Statistical significance was assessed using two-way ANOVA with Sidak’s multiple-comparisons test.

Because iron solubility increases under acidic conditions^44^, we tested whether pH modulates PEPR signaling under low Pi. Although alkaline pH enhances Pep1–PEPR binding *in vitro*^45^, Pep1-induced defense gene expression in our system was elevated under acidic conditions (Fig. 4c), consistent with increased iron availability promoting immune potentiation *in vivo*. However, Pep1-induced seedling growth inhibition was unaffected by pH conditions (Supplementary Fig. 6d), further supporting the conclusion that Pep-triggered immunity and growth regulation are not mutually dependent downstream outputs.

Under Pi deficiency, Fe³□ accumulation in the root meristem is strongly reduced in *lpr1 lpr2* mutants^43^. Consistently, enhanced Pep1-induced gene expression under low Pi remained reduced in *lpr1 lpr2* even in the presence of excess iron (Fig. 4d), indicating that LPR1/LPR2-mediated redox regulation is required for iron-dependent amplification of PEPR outputs.

Excess iron can elevate ROS and reinforce defense responses, while increasing oxidative stress ^17,46,47^. Under low Pi with excess iron, ROS accumulation increased in root tips (Supplementary Fig. 6e), as previously described^43^. In *rbohD rbohF* mutants defective in apoplastic ROS production ^48,49^, Pep1-induced defense gene expression was reduced under low Pi (Fig. 4e), demonstrating that ROS production is required for iron-dependent enhancement of PEPR signaling.

Finally, several defense regulators, including *FLAVIN-DEPENDENT MONOOXYGENASE1* (*FMO1*), *ENHANCED DISEASE SUSCEPTIBILITY5* (*EDS5*), and *PHYTOALEXIN DEFICIENT3* (*PAD3*)^50–52^, were induced by low Pi or excess iron alone, even without exogenous Pep1 application (Supplementary Fig. 6f). RNA-seq analysis confirmed that these genes were upregulated under Pi deficiency (Supplementary Fig. 6g, h). However, their induction was diminished in *pepr1 pepr2*, indicating that endogenous PEPR signaling contributes to iron-associated defense activation under Pi limitation.

### WRKY33 contributes to PEPR-dependent immune amplification under Pi limitation

*Cis*-regulatory motif enrichment analysis of Pep1-inducible genes whose expression was further enhanced under Pi deficiency revealed significant enrichment of WRKY33-binding motifs, including those recognized by its closest paralogue, WRKY25, at both 2 h and 10 h after Pep1 treatment (Fig. 5a). WRKY33 is a central regulator of camalexin biosynthesis and defense-associated transcriptional reprogramming^53–55^. Beyond immunity, WRKY33 has also been implicated in nutrient stress responses, including the response to Pi starvation, suggesting a potential role in integrating nutrition and defense signals^56–58^.

**Fig. 5:**
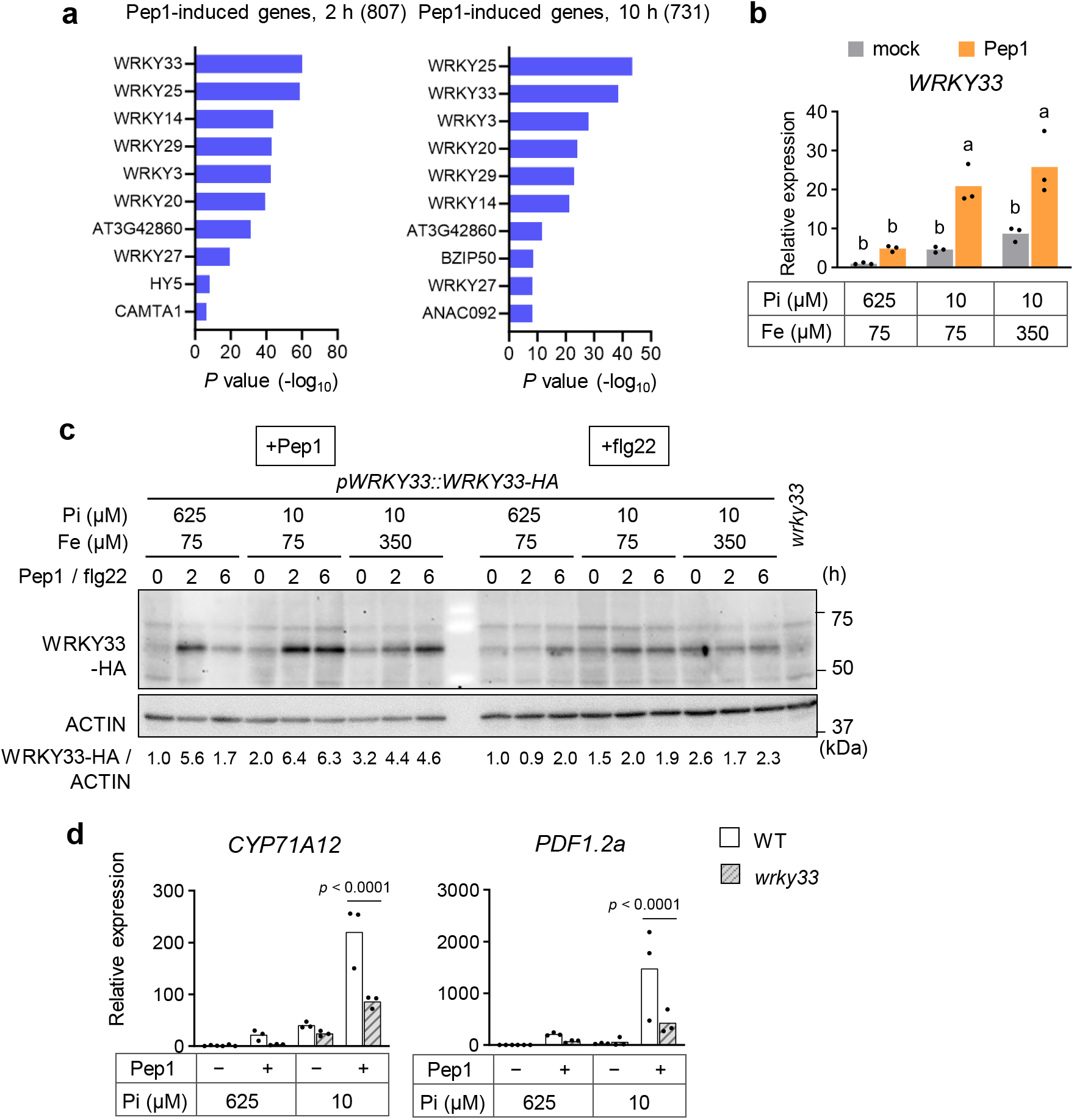
WRKY33 contributes to PEPR-dependent transcriptional amplification under Pi limitation. **a**, Top 10 enriched *cis*-regulatory elements among the Pep1-induced genes (log_2_FC > 1, FDR < 0.05). **b**, *WRKY33* transcript levels in WT roots grown under the indicated Pi and Fe conditions and treated with 1 µM Pep1 for 6 h. Data are means, *n* = 3 biological replicates. Different letters indicate significant differences (*p* < 0.05; two-way ANOVA followed by Tukey’s multiple-comparisons test). **c**, Immunoblot analysis of WRKY33-HA accumulation in the *pWRKY33::WRKY33-HA* expression line. Seedlings were grown under the indicated Pi and Fe conditions and treated with 1 µM Pep1 or 1 µM flg22 for the indicated times. ACTIN served as a loading control (bottom). WRKY33–HA band intensities normalized to ACTIN are shown, relative to the 625 µM Pi 0-h sample (set to 1) within each elicitor series. Similar results were obtained in three independent experiments. **d**, *CYP71A12* and *PDF1.2a* transcript levels in the WT and *wrky33* roots grown under Pi-sufficient conditions (+Pi, 625 µM) or Pi-deficient conditions (−Pi, 10 µM) and treated with 1 µM Pep1 for 6 h. Data are means, *n* = 3 biological replicates. *p* values for significant comparisons are indicated in the figure. Statistical significance was assessed using two-way ANOVA with Sidak’s multiple-comparisons test.

Consistent with this notion, *WRKY33* expression was induced by Pep1 under low Pi and excess iron conditions (Fig. 5b). Immunoblot analysis further showed that WRKY33 protein accumulation was induced by Pep1 and that this accumulation was further enhanced under low Pi and excess iron conditions. By contrast, flg22-induced WRKY33 accumulation was not enhanced under these conditions (Fig. 5c), linking increased WRKY33 accumulation to the selective potentiation of Pep signaling. Given the established role of WRKY33 in JA-responsive defense gene expression^59^, its enhanced accumulation likely contributes to the preferential reinforcement of JA-mediated transcriptional outputs during PEPR signaling. Indeed, Pep1 induction of *CYP71A12* and *PDF1.2a* expression under low Pi was reduced in *wrky33* mutants relative to WT (Fig. 5d), demonstrating that WRKY33 is required for the full activation of these PEPR-mediated transcriptional outputs during Pi limitation.

### PEPR1/PEPR2 promote PSR and shape root microbiota partly through PHR1/PHL1 under Pi limitation

RNA-seq analysis revealed that Pep1 application to Pi-deprived seedlings led to suppression of phosphate starvation–inducible (PSI) genes (Fig. 6a), extending the previously reported antagonism between acute PTI activation and PSR outputs to PEPR-mediated immunity^37,38^. Notably, however, in the absence of exogenous Pep1, PSI gene induction under Pi deficiency was reduced in *pepr1 pepr2* compared with WT (Fig. 6a). These suppressed genes included canonical PHR1/PHL1 targets involved in Pi uptake (Fig. 6b), indicating that basal PEPR signaling contributes positively to PSR-associated transcription in non-elicited plants. Although WT and *pepr1 pepr2* seedlings exhibited comparable shoot biomass under sterile low Pi conditions, constitutive expression of PEPR1-FLAG or PEPR2-FLAG in the *pepr1 pepr2* background significantly increased shoot biomass (Fig. 6c, d), indicating that enhanced PEPR activity can promote plant growth under Pi limitation. 16S rRNA amplicon sequencing analysis revealed shifts in root microbiota composition in *phr1 phl1*, *pepr1 pepr2*, and *lpr1 lpr2* compared with WT, in nutrient-deficient soil, indicated by PERMANOVA analysis (Fig. 6e; Supplementary Fig. 7a). Despite these community-level differences, the relative abundances of dominant bacterial genera were broadly similar between WT and *pepr1 pepr2*, and no individual ASVs or genera were significantly enriched or depleted after multiple-testing correction (Supplementary Fig. 7b–d). These results suggest that the differences reflect coordinated but individually modest changes across the community rather than robust shifts in particular taxa. Microbiota differences associated with *pepr1 pepr2* and *lpr1 lpr2* were partially attenuated in the *phr1 phl1* background, and PERMANOVA indicated weaker community separation when the *pepr1 pepr2* mutations were introduced into the *phr1 phl1* background (Supplementary Fig. 7a). These findings indicate that PEPR1 and PEPR2 influence root microbiota assembly under nutrient limitation, in a partly PHR1/PHL1-dependent manner.

**Fig. 6:**
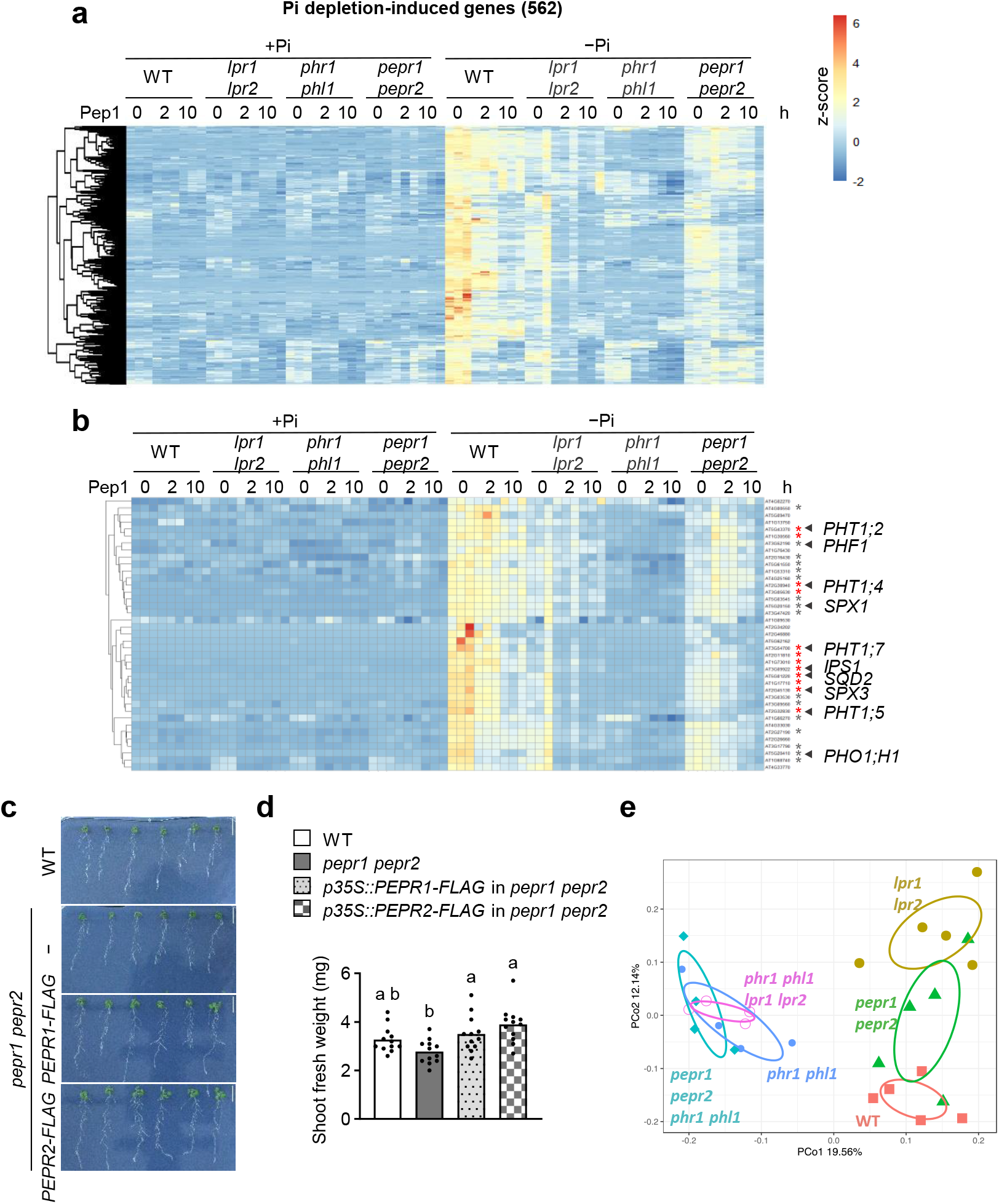
PEPR1/PEPR2 promote PSR and shape root microbiota under Pi limitation partly via PHR1/PHL1. **a**, Transcript profiles of the low Pi-induced gene cluster (log□FC > 1, FDR < 0.05). Log□(TPM) values were normalized as z-scores. **b**, Transcript profiles of representative low Pi-induced genes (log□FC > 1, FDR < 0.05). Log□(TPM) values were normalized as z-scores. Asterisks mark genes with higher expression in WT than in *pepr1 pepr2* under low Pi without exogenous Pep1 treatment (red, log□FC > 1, FDR < 0.05; gray, log□FC > 0.5, FDR < 0.05). **c**, Three-day-old seedlings were exposed to Pi-deficient conditions (10 µM Pi) for further 10 days. Scale bar, 20 mm. **d**, Shoot fresh weight of seedlings shown in c. Data are means, *n* = 12 biological replicates. Different letters indicate significant differences (*p* < 0.05; two-way ANOVA followed by Tukey’s multiple comparisons test). **e**, PCoA of the root microbiota of PSR regulator mutants and *pepr1 pepr2* grown in nutrient-deficient soil (*n* = 5, 5, 5, 4, 4, and 4 biological replicates for WT, *pepr1 pepr2*, *lpr1 lpr2*, *phr1 phl1*, *phr1 phl1 lpr1 lpr2*, and *phr1 phl1 pepr1 pepr2*, respectively).

### Non-canonical PROPEP-derived forms modulate PEPR signaling under Pi limitation

Although Pep-triggered PEPR activation typically inhibits plant growth^34,60^, our data indicate that basal PEPR activity promotes PSR transcription and that increased PEPR expression enhances biomass accumulation under low Pi. Immunoprecipitation–mass spectrometry (IP–MS) identified PROPEP6 in PEPR1 immunoprecipitates, with their association reduced following Pep1 treatment (Supplementary Fig. 8a, Supplementary Data 7). These results identify PROPEP6 as a PEPR1-associated protein under our experimental conditions. We therefore investigated whether PROPEP-derived forms modulate PEPR signaling. Transcripts of *PROPEP1*–*PROPEP8* were detected in non-elicited plants, with *PROPEP1/2/3/4* up-regulated under Pi deficiency relative to Pi sufficiency (Supplementary Fig. 9a, b), suggesting that multiple PROPEPs could contribute to basal PEPR regulation under low Pi.

Transient expression of Arabidopsis PROPEPs in *Nicotiana benthamiana* showed that several PROPEP members, including PROPEP6, suppressed Pep1-induced ROS production (Fig. 7a). We focused on PROPEP6 because, unlike the other family members, it did not yield a detectable product of the size expected for a mature Pep peptide (Supplementary Fig. 8b), consistent with its reported resistance to cleavage by METACASPASE4^61^. We also detected extracellular PROPEP6-derived forms in Arabidopsis, including a band corresponding to an apparently full-length PROPEP6–mCitrine fusion (∼39 kDa) (Supplementary Fig. 8c). The cleavage pattern differed from that observed in *N. benthamiana,* possibly reflecting species-specific proteolytic processing. Co-expression of PROPEP6 with PEPR1 in *N. benthamiana* did not induce a detectable increase in ROS production in the absence of Pep1 (Supplementary Fig. 8d). Co-treatment with recombinant PROPEP6 reduced ROS production in Pep1-treated WT *Arabidopsis* leaf discs, whereas PROPEP6 alone did not elicit a detectable increase under our assay conditions (Supplementary Fig. 8e). PROPEP6 had no detectable effect in *pepr1 pepr2* (Supplementary Fig. 8e). Co-treatment with recombinant PROPEP1 likewise reduced ROS production in Pep1-treated leaf discs, whereas PROPEP1 alone did not elicit a detectable increase (Supplementary Fig. 8f). Together with the agonist activity of mature Pep1, these findings suggest that different molecular forms of PROPEP1 can exert opposing effects on immune signaling.

**Fig. 7:**
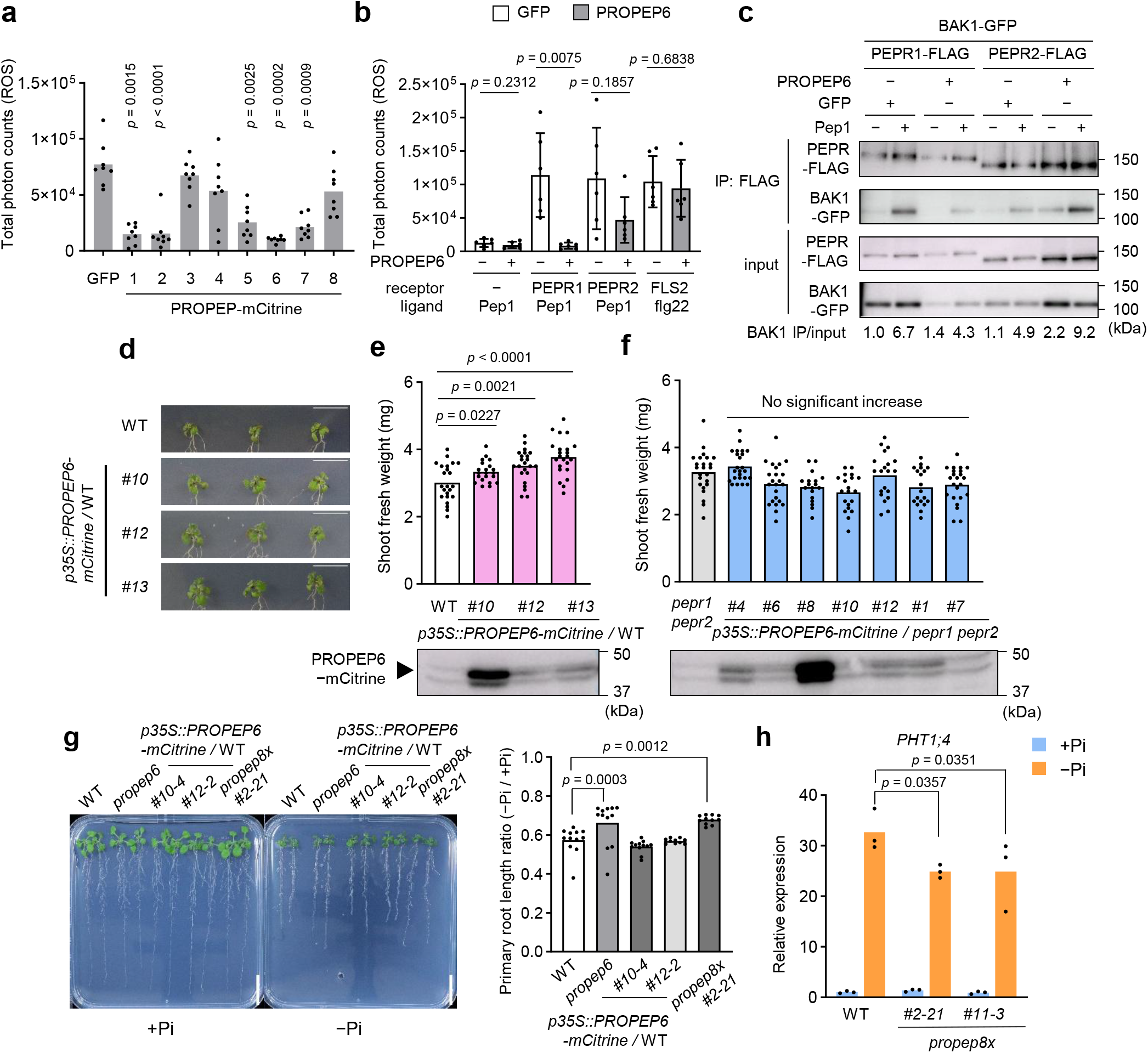
Non-canonical PROPEP-derived forms modulate PEPR signaling under phosphate limitation. **a**, Pep1-induced ROS production in *N. benthamiana* leaves co-expressing Arabidopsis PEPR1 and the indicated PROPEPs. **b**, ROS production in *N. benthamiana* leaves expressing the indicated receptors with untagged PROPEP6 or GFP and treated with Pep1 or flg22 as indicated. In **a**, data are means, *n* = 8 biological replicates. *p* values for significant comparisons are indicated. Statistical significance was assessed using one-way ANOVA with Dunnett’s multiple comparisons test against the GFP control. In **b**, data are means ± SD, *n* = 8 biological replicates. Statistical significance between ± PROPEP6 was determined using two-tailed t test. **c**, Co-immunoprecipitation analysis of PEPR1–BAK1 and PEPR2-BAK1 complex formation in *N. benthamiana* leaves co-expressing untagged PROPEP6 or GFP and treated with water or 1 µM Pep1 for 15 min. BAK1 was detected by immunoblotting with an anti-GFP antibody. Input proteins are shown. Similar results were obtained in three independent experiments. **d,e,** Representative seedlings (d) and shoot fresh weight (e) of WT and independent *p35S::PROPEP6-mCitrine* lines in the WT background. Three-day-old seedlings were exposed to Pi-deficient conditions (15 µM Pi) for further 10 days. Scale bar, 20 mm. Data are means, with *n* = 21 biological replicates for line *#10* and *n* = 23 for WT and the other lines. Statistical significance was assessed using two-sided student’s *t***-**test against WT. **f**, Shoot fresh weight of seedlings grown as in d,e. Data are means, with *n* = 24, 24, 23, 18, 20, 20, 19, and 23 biological replicates for *pepr1 pepr2*, line *#4*, *#6*, *#8*, *#10*, *#12*, *#7*, and *#1*, respectively. Statistical significance was assessed using two-sided student’s *t*-test against *pepr1 pepr2*. Immunoblots for PROPEP6-mCitrine accumulation are shown in the bottom (e,f). **g**, Primary root length of seedlings grown under +Pi (625 µM) or −Pi (50 µM). Scale bar, 20 mm. Data are means, with *n* = 12, 12, 12, 10, and 11 biological replicates for WT, *propep6*, line #*10-4*, #*12-2*, and *propep8x*, respectively. p values for significant comparisons are indicated. Statistical significance was assessed using one-way ANOVA with Dunnett’s multiple comparisons test against WT. **h**, *PHT1;4* transcript levels in roots of 13-day-old seedlings grown under +Pi or −Pi. Data are means, *n* = 3 biological replicates. *p* values for significant comparisons are indicated in the figure. Statistical significance between genotypes was assessed using two-way ANOVA with Tukey’s multiple comparisons test).

Notably, both PROPEP6–mCitrine and untagged PROPEP6 strongly suppressed Pep1-induced ROS production and MAPK activation when co-expressed with PEPR1, without affecting PEPR2-mediated signaling (Fig. 7a, b; Supplementary Fig. 8g). Unexpectedly, both PROPEP6 forms also reduced ROS production following flg22 treatment in PEPR1-expressing leaves (Supplementary Fig. 8h), whereas PROPEP6 did not significantly suppress this response in FLS2-expressing leaves (Fig. 7b). PEPR1/PEPR2 have been reported to associate with FLS2 and contribute to flg22-induced ROS production in *Arabidopsis*^62^. These observations raise the possibility that PROPEP6 attenuates flg22-triggered ROS production by suppressing the contribution of PEPR1 to FLS2 signaling.

Co-immunoprecipitation in *N. benthamiana* and *in vitro* binding assays further demonstrated physical association between PROPEP6 and PEPR1 (Supplementary Fig. 8i, j). PROPEP6 attenuated Pep1-induced PEPR1–BAK1 complex formation but did not affect PEPR2–BAK1 association (Fig. 7c). To further assess PROPEP6-mediated suppression of signaling, we examined MAPK phosphorylation in response to increasing Pep1 concentrations. PROPEP6 markedly suppressed MAPK phosphorylation induced by 1 µM Pep1, whereas 10 µM Pep1 restored MAPK phosphorylation to levels comparable to those observed in the absence of PROPEP6 (Supplementary Fig. 8k). These results indicate that PROPEP6-mediated suppression of Pep1 signaling can be overcome by high Pep1 concentrations. Suppression was also observed with untagged PROPEP6 (Fig. 7b, c, Supplementary Fig. 8h, and k), excluding that it was caused by the fluorescent protein fusion.

To assess the contribution of endogenous PROPEPs, we generated CRISPR/Cas9 mutants targeting *PROPEP1*, *PROPEP2*, *PROPEP3*, and *PROPEP6*. Higher-order *propep* mutants displayed enhanced Pep1-induced defense gene expression, particularly under low Pi (Supplementary Fig. 10a), supporting a role for endogenous PROPEPs in constraining PEPR-mediated immune outputs. The *propep6* single mutant showed no detectable enhancement of Pep1-induced defense gene expression (Supplementary Fig. 10b), suggesting partial redundancy among PROPEP family members in this context.

Given the positive role of PEPR1/PEPR2 in PSR transcription and growth under low Pi, we next examined the impact of PROPEP6 on these processes. Constitutive *PROPEP6-mCitrine* expression increased shoot biomass under low Pi in a *PEPR*-dependent manner (Fig. 7d–f), whereas loss of *PROPEP6* alone did not significantly affect shoot biomass (Supplementary Fig. 10c). By contrast, loss of *PROPEP6* alleviated primary-root growth inhibition under low Pi to a similar extent as the *propep* octuple mutant, indicating that PROPEP6 is the principal PROPEP contributing to this root-growth phenotype (Fig. 7g). Constitutive expression of *PROPEP6-mCitrine* did not further alter primary-root growth relative to WT under low Pi (Fig. 7g), suggesting that endogenous PROPEP6 is sufficient to exert this effect. Notably, *pepr1 pepr2* mutants exhibited an intermediate phenotype between WT and the *propep* mutants (Supplementary Fig. 10d), suggesting that PROPEP-dependent root-growth regulation is only partly mediated by PEPR1/PEPR2 and may involve additional signaling components. Consistent with these growth phenotypes, induction of *PHT1;4* under low Pi was reduced in the *propep* mutants (Fig. 7h). Collectively, these findings support a model in which non-canonical PROPEP-derived forms constrain Pep-PEPR1-mediated immunity while promoting PEPR-associated PSR.

## Discussion

Pi limitation imposes a fundamental challenge of balancing immune activation with nutrient acquisition. Our study identifies PEPR signaling as a context-responsive regulatory node within this balance, capable of biasing signaling toward defense or nutrient adaptation depending on ligand and nutritional context (Fig. 8). Quantitative proteomics and immunoblot analyses revealed that Pi deficiency differentially alters plasma-membrane accumulation of RKs and RLPs, maintaining or increasing PEPR1, PEPR2 and RLK7 while reducing several other PRRs, including FLS2 and CERK1 (Fig. 1). Beyond these changes in receptor abundance, selective regulation of downstream signaling makes a substantial contribution to the nutrient-dependent remodeling of immunity. Under Pi limitation, PEPR-mediated immunity is potentiated through selective amplification of particular outputs: PEPR–BAK1 complex formation and the acute Pep-triggered ROS burst remain largely unchanged, whereas MAPK activation and transcriptional outputs are enhanced, with WRKY33 contributing to the latter. Such uncoupling between ROS production and MAPK activation has been documented in PTI, in which these responses diverge downstream of PRRs and can be differentially regulated^63^. Pep1-induced resistance was detected more consistently in nutrient-deficient than in nutrient-sufficient soil, linking PEPR signaling to effective disease resistance under nutrient limitation (Fig. 2e; Supplementary Fig. 4). The relative contributions of Pi availability and other nutrient-dependent effects on host and pathogen physiology to this resistance phenotype remains to be resolved.

**Fig. 8:**
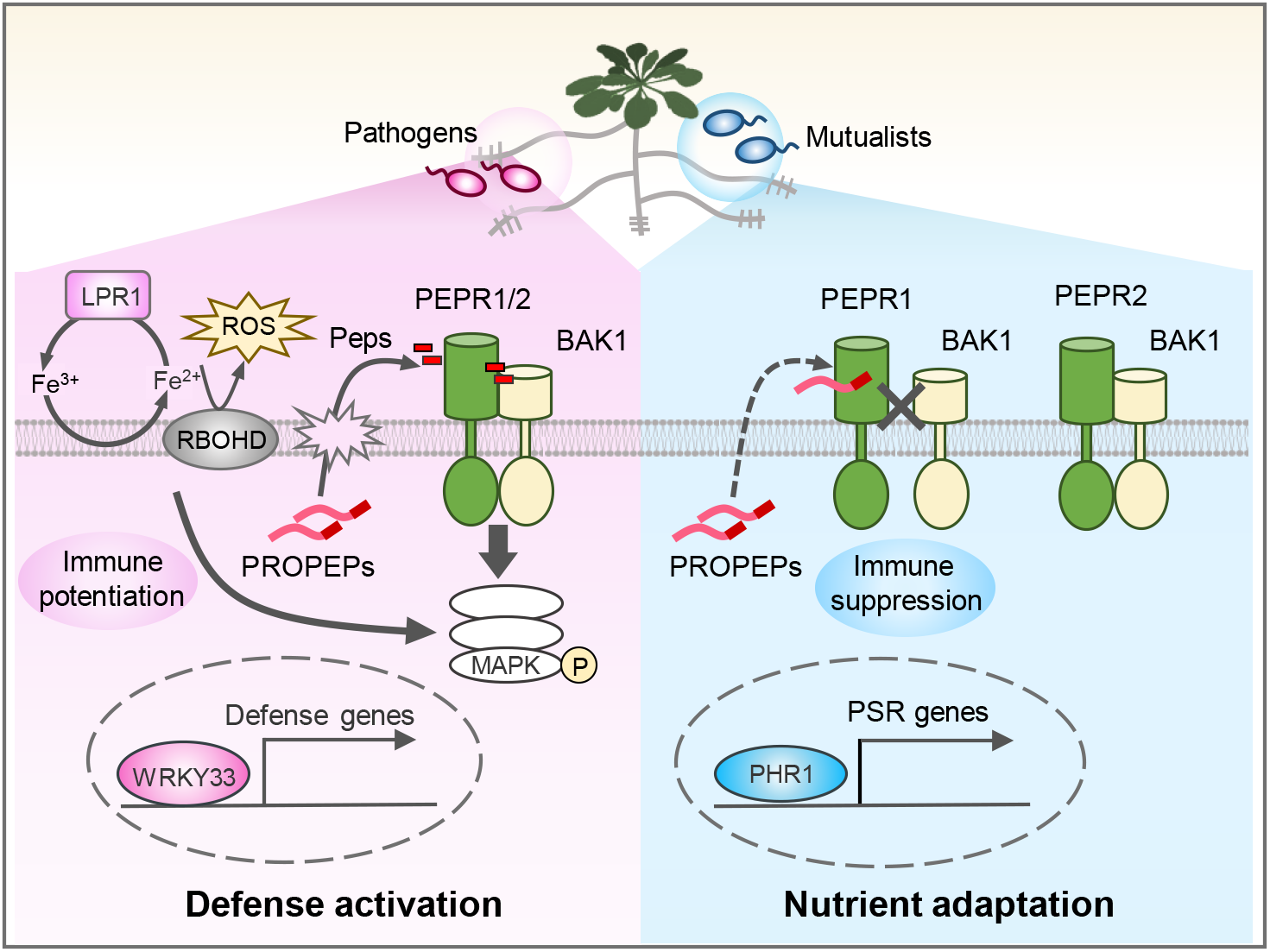
Model of iron- and PROPEP-dependent regulation of PEPR signaling during Pi limitation. Under low Pi, the abundance of PRRs is differentially regulated. Damage-associated Pep–PEPR signaling is selectively potentiated through the LPR1/LPR2- and RBOHD/RBOHF-dependent iron–redox module, involving iron accumulation, ROS production and WRKY33-mediated transcriptional amplification. When immunogenic Pep levels are low, PEPRs contribute to phosphate starvation responses (PSR). Distinct PROPEP-derived forms are proposed to preferentially constrain PEPR1-mediated immune signaling, while permitting PSR-associated growth. This context-dependent dual functionality likely enables PEPRs to balance defense and nutrient adaptation during Pi limitation.

The LPR1/LPR2–iron–ROS axis provides a mechanistic link between external Pi availability and PEPR-mediated immune potentiation. Excess iron enhances Pep-induced transcription and MAPK activation under low Pi, whereas iron deficiency attenuates these responses. This potentiation requires LPR1/LPR2 and RBOHD/RBOHF, implicating Pi- and iron-dependent redox remodeling despite the absence of a detectable increase in the acute Pep-triggered ROS burst. This distinction separates the broader redox regulation associated with Pi deficiency from the receptor-proximal ROS burst elicited by Pep. Potentiation of PEPR signaling is also associated with increased Pep-induced WRKY33 accumulation. By contrast, FLS2 signaling is attenuated under Pi deficiency, as previously described^39^, and is not potentiated by excess iron, while flg22-induced WRKY33 accumulation is not enhanced. Together, enrichment of WRKY33-binding motifs, selective enhancement of Pep1-induced WRKY33 accumulation, and the *wrky33* mutant phenotypes implicate WRKY33 in the selective amplification of PEPR-mediated transcriptional outputs under low Pi, although its mechanistic relationship to LPR1/LPR2-dependent iron–redox signaling remains unresolved. Differential connectivity of PRRs to membrane-associated signaling complexes or downstream signaling components may contribute to this specificity.

In the absence of acute elicitation by immunogenic Pep ligands, PEPR promotes PSR-associated transcription and growth under low Pi, indicating that basal PEPR activity engages nutrient-adaptation programs, including PHR1/PHL1-associated networks. Our findings further suggest that sensitization of the PEPR pathway is fine-tuned by non-canonical PROPEP-derived forms. Several PROPEPs suppress Pep1-induced responses upon transient expression, and recombinant PROPEP1 and PROPEP6 also suppress Pep1-induced ROS production upon acute co-treatment. Together with the established agonist activity of mature Peps, these findings support a model in which different PROPEP-derived forms modulate the activation threshold and output state of PEPR signaling.

PROPEP6 provides a model for this regulatory principle, exhibiting high resistance to proteolytic processing. Full-length PROPEP6 associates with PEPR1, attenuates Pep1-induced PEPR1–BAK1 complex formation, and suppresses PEPR1-mediated ROS production and MAPK activation in PEPR1-expressing leaves. Suppression can be overcome by increasing Pep1 concentrations, consistent with a model in which PROPEP6 modulates the threshold for PEPR1 signaling. The *propep6* single mutant did not exhibit enhanced Pep-triggered immune responses, whereas higher-order *propep* mutants did, indicating redundancy among PROPEP family members in constraining Pep-PEPR1-mediated immune signaling. By contrast, loss of PROPEP6 alone alleviated low Pi-induced primary root growth inhibition, suggesting that individual PROPEPs make distinct contributions to immune and PSR-associated PEPR outputs. The stronger root-growth phenotype of the *propep* mutants relative to *pepr1 pepr2* further suggests that PROPEP-dependent root-growth regulation is only partly mediated by PEPR1/PEPR2. Indeed, Pep7 is perceived by the LRR-RK SIRK1, which regulates aquaporin activity and lateral-root development with its coreceptor QSK1^64^. PROPEP-derived peptides may therefore regulate root development under Pi deficiency through a broader receptor network extending beyond PEPR1 and PEPR2.

The precise endogenous PROPEP-derived forms responsible for these activities remain to be defined. A limitation of this study is that the mechanisms underlying PROPEP-associated suppression remain unresolved and may differ among family members. The absence of a detectable ROS response to recombinant PROPEP1 or PROPEP6 alone and the recovery of MAPK activation at high Pep1 concentrations in the presence of PROPEP6 do not rule out constitutive low-level release of immunogenic Peps, which can cause receptor desensitization, as mechanisms contributing to the suppressive effects of some PROPEP family members. Likewise, it remains unclear whether basal PEPR-associated PSR reflects low-level constitutive signaling, ligand-independent receptor activity, or perception of alternative PROPEP-derived forms. Nevertheless, our genetic and biochemical data are compatible with the established model in which PROPEP processing generates immunogenic Pep ligands, while extending this model by suggesting that PROPEPs can also yield non-canonical forms that selectively modulate PEPR1 signaling, rather than serving solely as sources of immunogenic Pep ligands. Given the robust immune activation following exogenous Pep application and damage-induced Pep maturation through metacaspases^28,34,61,65^, we propose that the perception of immunogenic Pep ligands during damage or infection overcomes PROPEP-derived constraints, shifting the PEPR1 system from basal PSR-associated functions toward immune activation.

This model is consistent with an emerging view that endogenous peptide-receptor systems operate through ligand-context-dependent signal partitioning rather than simple on–off activation. In maize, ZmPep5a peptide competitively antagonizes immune responses induced by immunogenic ZmPeps^66^, and tomato antiSYS peptides function as antagonists required for immune homeostasis^67^. Distinct classes of RALF peptides can similarly direct FERONIA-associated signaling toward immune activation or suppression^68^. These examples support the possibility that structurally related peptide forms can produce opposing or complementary outputs within shared receptor networks.

The dual functionality of PEPR aligns with evidence that some endogenous peptide receptors regulate both immunity and development or nutrient responses. RLK7 perceives PIP peptides derived from PREPIP precursors to amplify immune signaling, while also regulating lateral root spacing through auxin-dependent pathways^69^. RLK7 additionally perceives CEP4 peptide, linking immune signaling with nitrogen starvation responses^70^. Similarly, PEPR signaling promotes root hair development^71,72^ in addition to its established immune functions. The maintenance of PEPR1 and RLK7 abundance under Pi deficiency raises the possibility that such multi-functional receptors help plants retain immune competence and developmental plasticity under nutrient limitation.

Our data establish iron as a key nutritional cue that potentiates PEPR-mediated immunity under Pi limitation. As an essential nutrient and catalyst of ROS generation, iron availability profoundly influences plant–microbe interactions^16,17,47^. Plants can restrict iron accessibility to pathogens or locally accumulate iron to enhance antimicrobial ROS production, whereas pathogens deploy siderophores to capture or scavenge host iron. Iron homeostasis and immune signaling are tightly interconnected: iron deficiency activates defense-associated pathways, while PTI signaling negatively regulates iron uptake programs^73–75^. A PROPEP2–PEPR2 module was recently shown to down-regulates iron uptake, rhizosphere acidification, and primary root growth during iron deficiency^75^. Together with our observation that Pep1-induced root growth inhibition occurs independently of iron availability (Supplementary Fig 5), these findings indicate that PEPR outputs differ in their dependence on iron-associated redox processes: iron is dispensable for Pep-induced growth inhibition but required for immune potentiation under low Pi. We propose that the LPR1/LPR2-dependent iron–redox module couples external Pi depletion to quantitative tuning of selective immune outputs downstream of PEPR.

PEPR activity under Pi limitation may also influence the root–microbiota interface. PSR alters root architecture, exudation patterns, and microbial community assembly^11,37,39,76^. We detected differences in overall microbial community composition between WT and *pepr1 pepr2*, although no individual ASVs or genera differed significantly after multiple-testing correction. The attenuation of these community differences in the *phr1 phl1* background suggests that PEPRs influence microbiota assembly partly through PHR1/PHL1-dependent nutrient-response networks. Thus, PEPR-dependent effects on the microbiota may arise primarily through changes in host nutritional physiology rather than direct activation of immunity.

Together, our results redefine PEPR signaling as a context-responsive receptor system with dual functions in immune activation and nutrient adaptation. Under Pi deficiency, increased PEPR abundance and LPR1/LPR2–dependent iron–ROS signaling potentiate selected immune outputs, with WRKY33 contributing to defense-related transcriptional amplification. In the absence of acute stimulation by immunogenic Pep ligands, basal PEPR activity instead supports PSR-associated functions, while non-canonical PROPEP-derived forms may help tune the balance between these functional states. Thus, rather than simply exemplifying an established nutrient–immunity trade-off, our findings reveal that the same receptor system can promote either immunity or nutrient-stress adaptation depending on ligand and nutritional context.

## Methods

### Plant materials and growth conditions

The *Arabidopsis thaliana* accession Columbia-0 (Col-0) was used as the wild type (WT) throughout this study. A complete list of plant materials is provided in Supplementary Data 8. Seeds were surface-sterilized in 5% sodium hypochlorite containing 0.05% Triton X-100 for 5 min, followed by three rinses with sterile distilled water. After stratification at 4°C for 2 days, seeds were sown on half-strength Murashige and Skoog (1/2 MS) medium containing 2.3 g/L MS basal salts (FUJIFILM Wako Pure Chemical), 1% (w/v) sucrose, and 1% (w/v) agar (Sigma-Aldrich, MO, USA). Plants were grown at 23°C under long-day conditions (16 h light/8 h dark) with 65% relative humidity. For phosphate deficiency assays, seedlings were initially cultivated on 1/2 MS medium for 4 days and then transferred either to control medium supplemented with 625 µM phosphate (+Pi) or to low-phosphate medium containing 10 µM phosphate (−Pi) for an additional 4 days. The +Pi medium consisted of 1/2 MS supplemented with 1% (w/v) sucrose and 1% (w/v) agarose (Biowest Regular Agarose G-10). For −Pi conditions, KH□PO□in the standard medium was replaced with KCl. The pH of both media was adjusted to 5.7 using 1 M KOH. For bacterial inoculation experiments, two-week-old seedlings were transplanted into soil and maintained in growth chambers at 22°C, 60% relative humidity, under short-day conditions (8 h light/16 h dark). Nutrient-deficient soil consisted of organic black soil (Maruishi Engei) mixed with Ezo sand (Kobayashi Sangyo) at a 2:1 (w/w) ratio. Plants were watered with distilled water. *N. benthamiana* plants were grown on 1/2 MS medium for 12 days under long-day conditions and subsequently transferred to the soil consisted of 1:1 (v/v) mixture of Super Mix A (Sakata Seed) and Vermiculite G20 (Maruishi Engei) Leaves from 4-to 5-week-old plants were used for transient expression assays.

### Plasmid DNA construction

The genomic region of *PROPEP*s, extending from the start codon to just before the stop codon, was amplified from wild-type Arabidopsis genomic DNA. PCR amplification was performed using PrimeSTAR GXL DNA Polymerase (Takara Bio, Cat# R050A). The amplified products were cloned into the pENTR/D-TOPO vector using the In-Fusion Snap Assembly Master Mix (Takara Bio, Cat# 638948). Proper insertion of the entry constructs was verified by Sanger sequencing with the BigDye Terminator v3.1 Cycle Sequencing Kit (Applied Biosystems, Cat# 4337455). Confirmed entry clones were then recombined into the destination vector pGWB502 (C-mCitrine) via the Gateway LR Clonase II Enzyme Mix (Invitrogen, Cat# 11791020), whereas entry clones containing the stop codon were transferred into the no-tag version of pGWB502 via the same Gateway LR reaction. To generate constructs for recombinant proteins for in vitro binding assays, PROPEP6 CDS was cloned into pCold GST DNA (Takara). The coding sequense of PEPR1, PEPR2, FLS2, and EFR extending from the start codon to just before the stop codon, and ectoPEPR1 (1-792 bp) were cloned into the pGWB502 (C-FLAG). The primer sequences used for cloning are listed in Supplementary Data 9.

### Generation of transgenic plants and *propep1/2/3/6* mutants

The p35S::PROPEP6-mCitrine construct described above were transformed into Col-0 plants by *Agrobacterium tumefaciens*-mediated transformation using the floral dip method^77^. For multiple sgRNA expression, complementary oligonucleotide pairs comprising the target sequence of *PROPEP*s (Supplementary Data 10) were annealed and then inserted into BsaI site and AarI site in pUC-AtU6-26_gtg (accession number: LC906469). For the construction of all-in-one CRISPR-Cas9 vectors, sgRNA expression cassettes from pUC-AtU6-26_gtg were transferred to pGHO-YC9 (accession number: LC906470).

### Plasma membrane enrichment

Plasma membrane fractions were enriched with minor modifications to a published protocol^78^. Briefly, whole 7-day-old Arabidopsis seedlings (∼200 mg fresh weight) grown under +Pi or −Pi conditions were frozen in liquid nitrogen, ground to a fine powder, and homogenized in 500 µl extraction buffer (250 mM sucrose, 50 mM Tris-HCl pH 7.5, 5% glycerol, 1 mM sodium molybdate, 25 mM NaF, 2 mM sodium orthovanadate, 10 mM EDTA, 3 mM DTT, and 1 mM PMSF). The homogenate was centrifuged at 8,000 × g for 10 min at 4°C, and the supernatant was transferred to ultracentrifuge tubes for subsequent fractionation according to the published protocol.

### Sample preparation for proteomic analysis

Proteins present in the supernatant were precipitated using pre-chilled acetone (−30°C), and the resulting pellets were resuspended in a digestion buffer containing 100 mM Tris-HCl (pH 9.0), 12 mM sodium lauryl sulfate (SLS), and 12 mM sodium deoxycholate (SDC). For each sample, 200 μg of protein was subjected to reduction and alkylation in a solution consisting of 10 mM TCEP, 40 mM chloroacetamide (CAA), and 50 mM ammonium bicarbonate for 30 min at 24°C in the dark. The samples were then diluted fivefold with 50 mM ammonium bicarbonate and digested overnight at 37°C with 2 μg of trypsin (Promega). Surfactants, including SLS and SDC, were removed using a phase-transfer method as previously described^79^. The resulting peptides were desalted using an in-house StageTip packed with SDB Empore disks (CDS), dried, and stored at −20°C until LC–MS/MS analysis.

### Sample preparation for IP-MS analysis

Arabidopsis seedlings were grown on either +Pi or −Pi medium for 7 days and then immediately frozen in liquid nitrogen. Seedlings were lysed with protein extraction buffer (50 mM pH7.5 Tris-HCl, 150 mM NaCl, 10% [v/v] glycerol, 0.5% [v/v] TritonX-100, supplemented with 1 mM EDTA (pH 8.0) and a protease inhibitor cocktail (Roche). Immunoprecipitation was performed using Anti-FLAG M2 Magnetic Beads (sigma), which were incubated with crude protein extracts for 1 h at 4°C. The beads were subsequently washed five times with protein extraction buffer. After the final wash, the beads were resuspended in 50 mM ammonium bicarbonate and transferred to a new microcentrifuge tube. An aliquot of the bead suspension was collected for immunoprecipitation (IP) samples. Bead-bound proteins were digested on-bead overnight at 37°C with trypsin (Promega). Following centrifugation, the peptide-containing supernatant was transferred to a fresh tube and subjected to reduction with 10 mM dithiothreitol (DTT) for 30 min at 25°C, followed by alkylation with 50 mM iodoacetamide (IAM) for 30 min at 25°C in the dark. The reaction was terminated by acidification with trifluoroacetic acid (TFA) to approximately pH 2. Peptides were desalted using C18 StageTips, eluted, and dried prior to LC–MS/MS analysis.

### LC-MS/MS analysis and raw data processing

Peptide samples were analyzed using an Easy-nLC 1200 nanoLC system (Thermo Scientific) coupled to an Orbitrap Exploris 480 mass spectrometer (Thermo Scientific) equipped with a FAIMS Pro interface. Dried peptides were reconstituted in 2% acetonitrile (ACN) with 0.1% formic acid (FA) and loaded onto a C18 capillary column (75 μm × 15 cm; Nikkyo Technos). Peptides were separated at 300 nL/min and 60°C using nonlinear gradients of 0.1% FA in water (Buffer A) and 0.1% FA in 80% ACN (Buffer B). A 140-min gradient was applied for global analyses, and a 105-min gradient for IP-MS samples. Eluted peptides were ionized at 2.2 kV and analyzed in positive mode using data-dependent acquisition. MS1 spectra (m/z 375–1500) were acquired at 120,000 resolution, and MS2 spectra (m/z >120) at 30,000 resolution. FAIMS compensation voltages of −40/−60 V or −50/−70 V were used under standard resolution settings (IT 100°C, OT 100°C). Peptide identification and MS1-based label-free quantification (LFQ) were performed with Proteome Discoverer 2.5 (Thermo Scientific) using the SEQUEST HT algorithm against the Araport11 database. Relative protein abundance shown in the figures corresponds to the scaled abundance values generated by Proteome Discoverer after software-based normalization (Scaling Mode). Search parameters included trypsin digestion (≤2 missed cleavages), precursor tolerance of 10 ppm, fragment tolerance of 0.02 Da, carbamidomethylation of cysteine as a fixed modification, and oxidation (M), N-terminal acetylation, and phosphorylation (S/T/Y) as variable modifications (≤3 per peptide). Peptide validation was conducted with Percolator, and only high-confidence identifications (FDR <1%) were used for protein inference and LFQ. Proteins identified based on these filtering criteria were used for downstream analyses. Total ion chromatograms were normalized across biological conditions. Missing quantitative values were retained as missing values and were not imputed prior to downstream analyses.

### qRT-PCR

Total RNA was isolated from root tissues using Sepasol-RNA I Super G (Nacalai Tesque). Following DNase treatment, cDNA was synthesized with the PrimeScript RT Reagent Kit with gDNA Eraser (Takara) according to the manufacturer’s protocol. qRT-PCR was conducted using Power SYBR Green PCR Master Mix (Applied Biosystems) on a Thermal Cycler Dice Real Time System III (Takara). Relative transcript levels were calculated by the ΔΔCt method and normalized to ACTIN2. Primer sequences are provided in Supplementary Data 9.

### RNA-sequencing and data analysis

Total RNA was extracted from plant root tissue, followed by DNase treatment using NucleoSpin RNA Plant (Macherey-Nagel, Cat# 740949.50) according to the manufacturer’s instruction. Each raw read data stored in DDBJ with accession numbers DRR to DRR was generated and analyzed as follows. Each RNA library was prepared from 1000 ng of 669 total RNA using the NEBNext Ultra II RNA Library Prep Kit for Illumina (New England 670 Biolabs, Cat# E7770) according to the manufacturer’s instruction. PCR for library amplification was performed for 7 cycles. The average length of the library was 350 bp, 672 as measured by a bioanalyzer. Concentrations were measured by Kapa Library. *Cis*-regulatory enrichment analyses were performed using ShinyGO 0.86 (https://bioinformatics.sdstate.edu/go/) with default settings.

### *Pto* DC3000 inoculation assay

*Pto* DC3000 was grown overnight at 28°C with shaking (180 rpm) in King’s B medium (2% Bacto Proteose Peptone No. 3, 0.15% K□HPO□, 1% glycerol, 5 mM MgSO□) supplemented with 50 μg/mL rifampicin. Bacterial cells were harvested and resuspended in sterile water to an OD□□□of 0.2. For induced-resistance assays, mature leaves were infiltrated with water, 1 µM Pep1 or 1 µM flg22 24 h before bacterial inoculation. For inoculation, the suspension was diluted 1 × 10□³ and infiltrated into three leaves per plant using a needleless syringe. Excess liquid was gently blotted from the leaf surface. Plants were maintained under standard growth conditions for the indicated times, with high humidity (∼95% RH) ensured by covering with a transparent plastic container. Bacterial populations were quantified 3 days after inoculation and expressed as cfu per mg fresh weight.

### ROS measurement

For total ROS measurements, 7-day-old seedlings were placed in 96-well plates containing 100 μL sterile water, covered, and incubated in the dark overnight. After 24 h, the water was replaced with 100 μL of assay solution containing 20 μM luminol L-012 (Sigma-Aldrich), 1 μg/mL horseradish peroxidase (HRP; Sigma-Aldrich), and 1 μM Pep1. Luminescence was recorded using a Fluoroskan Ascent FL microplate reader (Thermo Scientific), and ROS production was quantified as the cumulative photon counts.

For H2DCFDA staining, seedlings were grown on 1/2 MS medium for 4 days and then transferred to either +Pi or −Pi conditions for an additional 4 days. H2DCFDA was dissolved in DMSO (10 mg/mL), diluted to a 100× stock (1 mM) in 0.1 M phosphate buffer (pH 7.0), and stored at 4°C. Prior to use, aliquots were thawed in the dark, kept on ice, and diluted to 1× in phosphate buffer. Whole seedlings were incubated in 2 mL staining solution per well (12-well plate) for 10 min at room temperature in the dark. Fluorescence images were acquired with FV1000 confocal laser scanning biological microscope using a 10×/0.4 NA objective. H2DCFDA was excited at 488 nm, and emission was detected between 500–550 nm.

### Western blotting analysis

Frozen plant tissues were ground to a fine powder and homogenized in protein extraction buffer containing 62.5 mM Tris-HCl (pH 6.8), 10% glycerol, 2% SDS, 5% (v/v) 2-mercaptoethanol, and 0.02% bromophenol blue, supplemented with 1 mM EDTA (pH 8.0), phosphatase inhibitors, and a protease inhibitor cocktail (Roche). The buffer was added at a ratio of 3 μL per mg fresh weight. Samples were heated at 95°C for 5 min and centrifuged at 20,000 × g for 10 min at 25°C. The supernatants were collected for subsequent analysis. Proteins were resolved on 10% polyacrylamide gels and transferred onto PVDF membranes (Millipore). Membranes were blocked with 2% (w/v) skim milk (Nacalai Tesque), in TBS-T (25 mM Tris, 192 mM glycine, 0.1% [v/v] Tween-20) for 1 h at room temperature with agitation. After blocking, membranes were briefly rinsed with deionized water and TBS-T, followed by overnight incubation at 4°C with primary antibodies in TBS-T. Membranes were washed five times with deionized water and once in TBS-T for 5 min at room temperature, then incubated with secondary antibodies for 1 h at room temperature with agitation. After secondary antibody incubation, membranes were washed as described above. Signals were detected using Chemi-Lumi One L (Nacalai Tesque) and visualized with a FUSION FX imaging system (Vilber). The following antibodies were used: anti-pERK antibody (Cell Signaling, Cat#4370, dilution 1:5,000), anti-PROPEP3^36^ (rabbit polyclonal antibody, dilution 1:5,000), anti-BAK1^68^ (rabbit polyclonal antibody, dilution 1:5,000), anti-GFP (MBL, Cat#598, dilution 1:5000), anti-FLAG (Sigma-Aldrich, Cat# F1804, dilution 1:5000), HRP-conjugated anti-HA (Roche, Cat# 12013819001, dilution 1:5000), anti-ACTIN (Sigma-Aldrich, Cat# SAB4301137-100UL, dilution 1:5000), anti-RPN6 (Agrisera, Cat# AS15 2832A, dilution 1:5000), HRP-conjugated anti-rabbit IgG (Cell Signaling Technology, Cat# 7074, dilution 1:5000) and HRP-conjugated anti-mouse IgG (Cell Signaling Technology, Cat# 7076, dilution 1:5000). All antibodies were diluted in TBS-T.

### Co-immunoprecipitation (Co-IP) assays

Arabidopsis seedlings were grown on either +Pi or −Pi medium for 7 days, treated with 1 μM Pep1 for 15 min, and then immediately frozen in liquid nitrogen. Roots were lysed with protein extraction buffer (50 mM pH7.5 Tris-HCl, 150 mM NaCl, 10% [v/v] glycerol, 0.5% [v/v] TritonX-100, supplemented with 1 mM EDTA (pH 8.0) and a protease inhibitor cocktail (Roche). Immunoprecipitation was performed using Anti-FLAG M2 Magnetic Beads (sigma), which were incubated with crude protein extracts for 1 h at 4°C. The beads were subsequently washed five times with protein extraction buffer, resuspended in SDS sample buffer, and incubated at 95°C for 5 min. Proteins were analyzed by SDS-PAGE followed by immunoblotting.

### Extracellular PROPEP detection assays

Seven-day-old seedlings grown under either +Pi or −Pi conditions were transferred to liquid MS medium. After 24 h, the medium was replaced with fresh medium prior to Pep1 treatment. The following day, seedlings were treated with 1 µM Pep1 for 24 h. Culture media were collected, and proteins were concentrated using Strataclean resin (Agilent Technologies). After centrifugation, the resin was boiled in SDS sample buffer, and proteins were separated by SDS–PAGE. Seedling tissues were also harvested and subjected to immunoblot analysis.

### DNA extraction and 16S rRNA gene amplicon sequencing

Root tissues were homogenized using a Mixer Mill (Verder Scientific Co., Ltd., Tokyo, Japan), and total genomic DNA was isolated with the NucleoSpin® Soil kit (MACHEREY-NAGEL GmbH & Co. KG, Germany) in accordance with the manufacturer’s protocol. The V4 region of the 16S rRNA gene was amplified by touchdown PCR using KOD FX Neo polymerase (TOYOBO Co., Ltd., Osaka, Japan) together with Illumina adapter-linked barcoded primers 515F (TCGTCGGCAGCGTCAGATGTGTATAAGAGACAGGTGCCAGCMGCCGCGGTA-A) and 806R (GTCTCGTGGGCTCGGAGATGTGTATAAGAGACAGGGACTACHVGG-GTWTCTAAT), as previously described^80^. PCR products of the expected size were separated on a 1% agarose gel, excised, and purified using the FastGene Gel/PCR Extraction Kit (Nippon Genetics Co., Ltd., Tokyo, Japan) according to the manufacturer’s instructions. Amplicon libraries were subsequently constructed using Nextera XT Index kits (Illumina, Inc., CA, USA), and 300-bp paired-end sequencing was carried out on an Illumina MiSeq platform with the MiSeq Reagent Kit v3.

### Sequence data processing

Raw sequencing reads were demultiplexed using the clsplitseq command implemented in Claident^81^, applying a minimum quality threshold of 30. Amplicon sequence variants (ASVs) were inferred using the DADA2 pipeline^82^ in the R package dada2. For the filterAndTrim() function, parameters were specified as follows: truncQ = 2; truncLen = 220 (forward) and 150 (reverse); trimLeft = 19 (forward) and 20 (reverse); maxN = 0; and maxEE = 2 for both forward and reverse reads. Chimeric sequences were detected and eliminated using default settings. Taxonomic assignment of ASVs was conducted with SINA v1.2.11^83^ against the SILVA SSURef NR99 database release 132^84^, applying a minimum similarity cutoff of 85%. Sequences classified as Eukarya, Archaea, mitochondrial or chloroplast in origin, as well as those that could not be aligned to reference sequences, were excluded from further analyses.

### Statistical analyses

Beta diversity analyses were performed on the Galaxy Europe platform using the QIIME2 wrapper developed by the European Galaxy Team. Bray–Curtis dissimilarity matrices were generated, and statistical differences among groups were evaluated by PERMANOVA (beta-group-significance, method = permanova) with pairwise comparisons and 999 permutations. Differences between groups were determined based on the separation of multivariate centroids in the distance matrix space. Statistical significance was assessed using false discovery rate (FDR)-adjusted q-values provided by Galaxy. Data visualization was carried out in R using the ggplot2 package^85^.

### Agrobacterium-mediated transient expression

Agrobacterium strains (GV3101::pMP90) carrying the relevant expression vectors were cultured overnight at 28°C with shaking (180 rpm) in 2×YT liquid medium [1% (w/v) yeast extract, 1,6% (w/v) tryptone, 0.5% (w/v) NaCl] supplemented with 50 µg/mL gentamycin and 100 µg/mL spectinomycin. Bacterial cells were collected by centrifugation at 2,000 g for 10 min at 25°C, washed once with sterilized water, and resuspended in sterilized water to an OD600 of 0.5. Acetosyringone was added to the suspension to a final concentration of 500 µM. The bacterial suspensions were infiltrated into the leaves of *N. benthaminana* using a needleless syringe The infiltrated plants were maintained under their growth conditions for approximately 48 hours.

### In vitro protein interaction assays

GST-tagged recombinant proteins were expressed in *E. coli* BL21 and purified using Glutathione Sepharose 4B (GE Healthcare) according to the manufacturer’s instructions. Beads were washed with wash buffer (50 mM Tris-HCl, pH 8.0, 100 mM NaCl), and proteins were eluted with elution buffer (50 mM Tris-HCl, pH 8.0, 10 mM reduced glutathione). Protein concentrations were determined by the Bradford assay. Expression constructs encoding ectoPEPR1-FLAG and GFP-FLAG were transformed into *A. tumefaciens* GV3101 and transiently expressed in *N. benthamiana* leaves. Two days after infiltration, leaf tissue was homogenized in extraction buffer (50 mM Tris-HCl, pH 7.5, 150 mM NaCl, 10% [v/v] glycerol, 0.5% [v/v] Triton X-100) supplemented with 1 mM EDTA (pH 8.0) and protease inhibitor cocktail (Roche). Extracts were incubated with anti-FLAG M2 magnetic beads (Sigma) for 1 h at 4°C, washed with extraction buffer, and eluted with TBS containing 500 μg/mL FLAG (DYKDDDDK) peptide. For pull-down assays, 2 μL of eluted ectoPEPR1-FLAG or GFP-FLAG were incubated with 150 nM purified GST-PROPEP6 in pull-down buffer (50 mM Tris-HCl, pH 7.5, 150 mM NaCl, 10% glycerol) for 2 h at 4°C. GFP-FLAG was used as control. Glutathione beads were then added and incubated for 2 h at 4°C. Beads were washed with buffer and boiled in SDS sample buffer prior to analysis.

## Data availability

The raw RNA-sequencing data have been deposited in the DDJB database under accession numbers PRJDB37679 and PRJDB35626.

LC-MS/MS raw data have been deposited in Japan Proteome Standard Repository/ Database (jPOSTP) under accession code PXD074930. The data can be accessed using the access key 7188.

The 16S rRNA gene amplicon and metagenome sequencing data have been deposited in the DNA Data Bank of Japan Sequence Read Archive (DRA) under the accession number DRA012727.

Source Data are provided with this paper.

## Acknowledgements

We thank Mie Matsubara and John Jewish Dominguez (Nara Institute of Science and Technology) for technical assistance; Jian-Feng Li (Sun Yat-sen University) for *propep8x* seeds^86^. This work was supported in part by JSPS KAKENHI Grant Numbers 25KJ1825 (N.T.), 18H02467 and 21H02507 (Y.S.).

## Author contributions

Y.S. conceived the study. N.T., T.H.L., K.H., and Y.S. designed the experiments. N.T., T.H.L., M.L., K.Y., K.H., T.H., S.Y., Y.D.U., M.F., M.F.C., and T.U. performed the experiments and analyzed the data. H.A. developed the materials. K.Y., Y.D.U., K.O., S.C., M.F.C., and T.U. advised on the data analyses. N.T. and Y.S. wrote the manuscript with contributions from the other authors.

