## Supplementary Information for "Iron-dependent reprogramming of damage-associated peptide receptor signaling coordinates immunity and phosphate stress adaptation"

**Supplementary materials**

Supplementary Figures 1-10

Supplementary Data 1-10

**Supplementary Data 1**

All identified protein in the quantitative proteomic analysis with plasma membrane enriched faction.

**Supplementary Data 2**

Differential gene expression analysis of root transcriptomes following Pep1 or chitin treatment under Pi-sufficient (+Pi) and Pi-deficient (−Pi) conditions.

**Supplementary Data 3**

Normalized expression values of Arabidopsis roots following Pep1 or chitin treatment under Pi-sufficient (+Pi) and Pi-deficient (−Pi) conditions obtained by RNA-seq.

**Supplementary Data 4**

Pep1-responsive genes ranked by the change in Pep1 responsiveness under Pi-deficient conditions (Δlog2FC).

**Supplementary Data 5**

Differential gene expression analysis of root transcriptomes from Arabidopsis WT, *phr1 phl1*, *lpr1 lpr2*, and *pepr1 pepr2* following 2 h or 10 h Pep1 treatment under Pi-sufficient (+Pi) and Pi-deficient (−Pi) conditions.

**Supplementary Data 6**

Normalized expression values of Arabidopsis WT, *phr1 phl1*, *lpr1 lpr2*, and *pepr1 pepr2* roots treated with Pep1 for 2 h or 10 h under Pi-sufficient (+Pi) and Pi-deficient (−Pi) conditions.

**Supplementary Data 7**

Complete list of proteins identified in the PEPR1 co-immunoprecipitation–mass spectrometry analysis

**Supplementary Data 8**

List of Arabidopsis plants used in this study.

**Supplementary Data 9**

List of primers used in this study.

**Supplementary Data 10**

List of target sequence of PROPEPs used for generation of *propep1/2/3/6*.

row min — row max

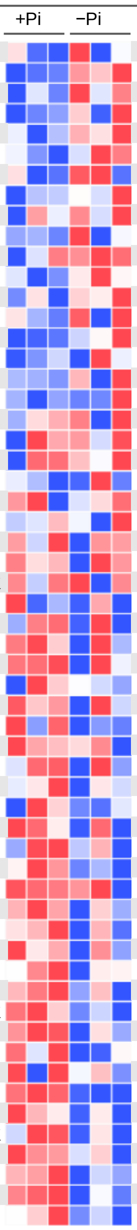

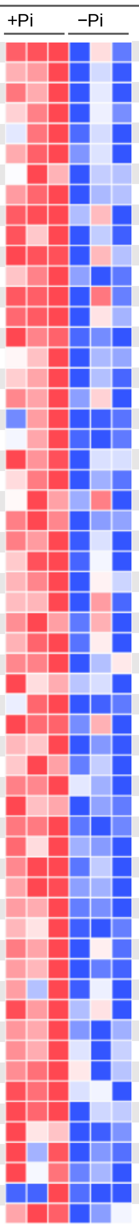

| Accession | Gene symbol | Protein family | Fold change (−Pi / +Pi) | *P*-value |
|---|---|---|---|---|
| AT1G58190.2 | RLP9 | RLP | 2.16 | 0.2547 |
| AT4G23190.1 | CRK11 | CRR | 1.88 | 0.0143 |
| AT1G75640.1 | MUS | LRR-RLK | 1.68 | 0.1953 |
| AT3G02130.1 | RPK2 | LRR-RLK | 1.56 | 0.2530 |
| AT1G51800.1 | IOS1 | LRR-RLK | 1.53 | 0.0575 |
| AT1G73080.1 | PEPR1 | LRR-RLK | 1.51 | 0.2141 |
| AT5G51350.1 | MOL1 | LRR-RLK | 1.41 | 0.2613 |
| AT2G01210.1 | ZAR1 | LRR-RLK | 1.41 | 0.1396 |
| AT3G21340.1 | | LRR-RLK | 1.31 | 0.4476 |
| AT1G60800.1 | NIK3 | LRR-RLK | 1.28 | 0.4602 |
| AT1G56140.1 | | LRR-RLK | 1.26 | 0.1675 |
| AT5G65240.2 | | LRR-RLK | 1.25 | 0.1994 |
| AT5G25930.1 | HSL3 | LRR-RLK | 1.23 | 0.2468 |
| AT5G59700.1 | ANJ | CrRLK1L | 1.22 | 0.4410 |
| AT1G51850.2 | SIF2 | LRR-RLK | 1.19 | 0.0681 |
| AT1G09970.2 | RLK7 | LRR-RLK | 1.17 | 0.3781 |
| AT1G28440.1 | HSL1 | LRR-RLK | 1.17 | 0.4244 |
| AT5G40380.1 | CRK42 | CRR | 1.06 | 0.4399 |
| AT3G13380.1 | BRL3 | LRR-RLK | 1.06 | 0.8289 |
| AT3G56100.1 | IMK3 | LRR-RLK | 1.04 | 0.7681 |
| AT4G23180.1 | CRK10 | CRR | 1.03 | 0.7365 |
| AT4G33430.2 | BAK1 | LRR-RLK | 1.03 | 0.6957 |
| AT1G53440.1 | | LRR-RLK | 1.02 | 0.9272 |
| AT2G33050.1 | RLP26 | RLP | 1.00 | 0.9831 |
| AT5G48380.1 | BIR1 | LRR-RLK | 0.96 | 0.6225 |
| AT5G40170.1 | RLP54 | RLP | 0.96 | 0.7832 |
| AT4G21390.1 | | G-type lecRLK | 0.94 | 0.6707 |
| AT3G14350.1 | SRF7 | LRR-RLK | 0.93 | 0.7270 |
| AT1G53430.1 | NILR2 | LRR-RLK | 0.92 | 0.4029 |
| AT1G70530.1 | CRK3 | CRR | 0.92 | 0.2534 |
| AT2G23770.1 | LYK4 | lysm-RLK | 0.90 | 0.2709 |
| AT1G70520.1 | CRK2 | CRR | 0.90 | 0.6611 |
| AT2G27060.1 | | LRR-RLK | 0.89 | 0.4696 |
| AT3G51740.1 | IMK2 | LRR-RLK | 0.89 | 0.1841 |
| AT5G60270.1 | | L-type lecRLK | 0.89 | 0.2666 |
| AT1G15530.1 | | L-type lecRLK | 0.88 | 0.2800 |
| AT4G08850.1 | MIK2 | LRR-RLK | 0.88 | 0.1935 |
| AT4G23270.3 | CRK19 | CRR | 0.88 | 0.4172 |
| AT4G18760.1 | RLP51 | RLP | 0.87 | 0.2139 |
| AT1G51805.1 | SIF3 | LRR-RLK | 0.87 | 0.3403 |
| AT2G16250.1 | | LRR-RLK | 0.86 | 0.0346 |
| AT3G56370.1 | IRK | LRR-RLK | 0.86 | 0.3858 |
| AT5G49760.1 | CARD1 | LRR-RLK | 0.85 | 0.0338 |
| AT3G14840.2 | LIK1 | LRR-RLK | 0.84 | 0.2060 |
| AT5G60300.3 | | L-type lecRLK | 0.84 | 0.3629 |
| AT1G66150.1 | TMK1 | LRR-RLK | 0.84 | 0.0278 |
| AT4G23250.2 | CRK17 | CRR | 0.83 | 0.1459 |
| AT1G11410.4 | | G-type lecRLK | 0.82 | 0.0025 |
| AT3G28450.1 | BIR2 | LRR-RLK | 0.81 | 0.0314 |
| AT1G68400.1 | | LRR-RLK | 0.81 | 0.0627 |
| AT4G21410.3 | CRK29 | CRR | 0.81 | 0.6743 |
| AT3G21630.1 | CERK1 | lysm-RLK | 0.80 | 0.0026 |
| AT1G10850.1 | | LRR-RLK | 0.80 | 0.1449 |
| AT1G11330.2 | | G-type lecRLK | 0.79 | 0.0919 |
| AT1G71830.1 | SERK1 | LRR-RLK | 0.79 | 0.1001 |
| AT4G39400.1 | BRI1 | LRR-RLK | 0.78 | 0.0351 |
| AT1G06840.1 | HAK1 | LRR-RLK | 0.77 | 0.0360 |
| AT2G37710.1 | | L-type lecRLK | 0.77 | 0.1529 |

| Accession | Gene symbol | Protein family | Fold change (−Pi / +Pi) | *P*-value |
|---|---|---|---|---|
| AT5G65700.2 | BAM1 | LRR-RLK | 0.76 | 0.0545 |
| AT3G02880.1 | QSK1 | LRR-RLK | 0.75 | 0.0573 |
| AT2G17120.1 | LYM2 | lysm-protein | 0.75 | 0.0799 |
| AT5G16590.1 | QSK2 | LRR-RLK | 0.75 | 0.0408 |
| AT2G02780.1 | | LRR-RLK | 0.74 | 0.0872 |
| AT3G51550.1 | FER | CrRLK1L | 0.74 | 0.0490 |
| AT1G30570.1 | HERK2 | CrRLK1L | 0.73 | 0.0209 |
| AT3G46290.1 | HERK1 | CrRLK1L | 0.73 | 0.0039 |
| AT5G38990.1 | MDS1 | CrRLK1L | 0.72 | 0.0855 |
| AT2G31880.1 | SOBIR1 | LRR-RLK | 0.72 | 0.0798 |
| AT3G24550.1 | PERK1 | PERK | 0.71 | 0.0899 |
| AT4G36180.1 | MUL | LRR-RLK | 0.71 | 0.0378 |
| AT1G53730.2 | SRF6 | LRR-RLK | 0.71 | 0.1546 |
| AT2G26730.1 | | LRR-RLK | 0.70 | 0.0512 |
| AT4G17090.1 | BAM3 | LRR-RLK | 0.70 | 0.0102 |
| AT5G10020.1 | SIRK1 | LRR-RLK | 0.69 | 0.0385 |
| AT3G08680.1 | | LRR-RLK | 0.69 | 0.0467 |
| AT1G07650.2 | LMK1 | LRR-RLK | 0.69 | 0.1486 |
| AT2G02220.1 | PSKR1 | LRR-RLK | 0.68 | 0.1190 |
| AT1G28340.1 | RLP4 | RLP | 0.67 | 0.0383 |
| AT2G41820.1 | PXC3 | LRR-RLK | 0.66 | 0.0701 |
| AT3G23750.1 | BARK1 | LRR-RLK | 0.66 | 0.0212 |
| AT4G22130.1 | SRF8 | LRR-RLK | 0.66 | 0.1719 |
| AT1G14390.1 | | LRR-RLK | 0.66 | 0.0101 |
| AT2G37050.3 | SIMP1 | LRR-RLK | 0.66 | 0.0624 |
| AT3G17840.1 | RLK902 | LRR-RLK | 0.65 | 0.0449 |
| AT5G58300.3 | KOIN | LRR-RLK | 0.64 | 0.0486 |
| AT1G12460.1 | | LRR-RLK | 0.62 | 0.2294 |
| AT4G23740.1 | | LRR-RLK | 0.62 | 0.0600 |
| AT2G33580.1 | LYK5 | lysm-RLK | 0.62 | 0.0679 |
| AT5G46330.2 | FLS2 | LRR-RLK | 0.62 | 0.0360 |
| AT3G28040.1 | | LRR-RLK | 0.62 | 0.0900 |
| AT4G32300.1 | | G-type lecRLK | 0.62 | 0.0362 |
| AT5G10290.2 | | LRR-RLK | 0.62 | 0.1072 |
| AT2G01820.1 | TMK3 | LRR-RLK | 0.61 | 0.0439 |
| AT4G20940.1 | GHR1 | LRR-RLK | 0.60 | 0.0103 |
| AT2G42800.1 | RLP29 | RLP | 0.59 | 0.0657 |
| AT1G56145.2 | CORK1 | LRR-RLK | 0.59 | 0.0522 |
| AT5G54380.1 | THE1 | CrRLK1L | 0.57 | 0.0013 |
| AT5G14210.2 | | LRR-RLK | 0.57 | 0.0622 |
| AT1G48480.1 | RKL1 | LRR-RLK | 0.56 | 0.0180 |
| AT3G57830.1 | | LRR-RLK | 0.55 | 0.0304 |
| AT1G27190.1 | BIR3 | LRR-RLK | 0.54 | 0.0328 |
| AT2G19230.1 | | LRR-RLK | 0.54 | 0.0154 |
| AT1G51940.1 | LYK3 | lysm-RLK | 0.52 | 0.1019 |
| AT5G63410.1 | | LRR-RLK | 0.50 | 0.0202 |
| AT4G02420.1 | | L-type lecRLK | 0.49 | 0.2621 |
| AT4G00340.1 | | G-type lecRLK | 0.49 | 0.1172 |
| AT4G11530.1 | CRK34 | CRR | 0.48 | 0.0048 |
| AT3G49670.1 | BAM2 | LRR-RLK | 0.47 | 0.0294 |
| AT4G04960.1 | | L-type lecRLK | 0.45 | 0.0858 |
| AT4G34220.1 | | LRR-RLK | 0.44 | 0.0622 |
| AT4G03390.2 | SRF3 | LRR-RLK | 0.43 | 0.0342 |
| AT5G63710.3 | | LRR-RLK | 0.41 | 0.0539 |
| AT1G21880.2 | LYM1 | lysm-protein | 0.39 | 0.1529 |
| AT1G75820.1 | CLV1 | LRR-RLK | 0.37 | 0.1040 |
| AT5G49660.1 | XIP1 | LRR-RLK | 0.36 | 0.4141 |
| AT1G63430.2 | | LRR-RLK | 0.34 | 0.0226 |

**Supplementary Fig. 1: Relative abundance of receptor kinases and receptor-like proteins under different Pi conditions.**

Relative protein abundance represents the scaled abundance values calculated by Proteome Discoverer. For visualization, abundance values were further scaled to the same range for each protein in the heatmap ($n$ = 3 biological replicates). Plants were grown under Pi-sufficient (+Pi, 625 µM) or Pi-deficient (−Pi, 10 µM) conditions. $P$ values were determined using a two-sided Student's $t$-test.

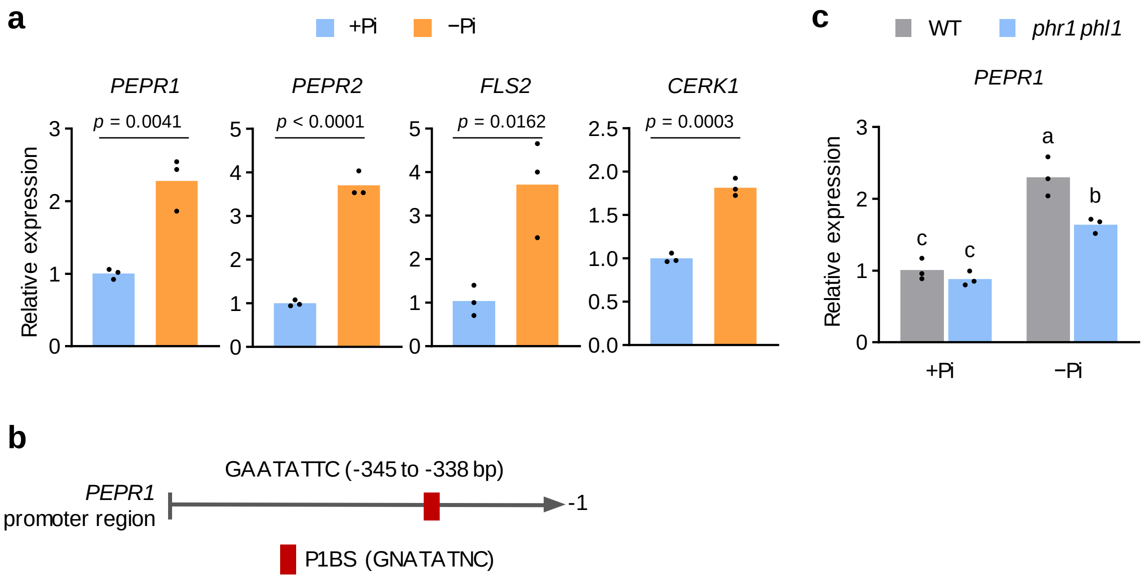


**Supplementary Fig. 2: Selective receptor abundance remodeling under low Pi is not explained by transcript level changes.**

**a**, Transcript levels of *PEPR1*, *PEPR2*, *FLS2* and *CERK1* in the WT roots grown under Pi-sufficient conditions (+Pi, 625 µM) or Pi-deficient conditions (−Pi, 10 µM). Data represent means, $n$ = 3 biological replicates. $P$-values were calculated using two-sided Student's $t$-test. **b**, Predicted PHR1-binding site (P1BS; GNATATNC) within the 931-bp *PEPR1* promoter region. **c**, Transcript levels of *PEPR1* in WT and *phr1 phl1* roots grown under +Pi and −Pi conditions. Data are means, $n$ = 3 biological replicates. Different letters indicate significant difference ($p$ < 0.05; two-way ANOVA followed by Tukey's multiple comparisons test).

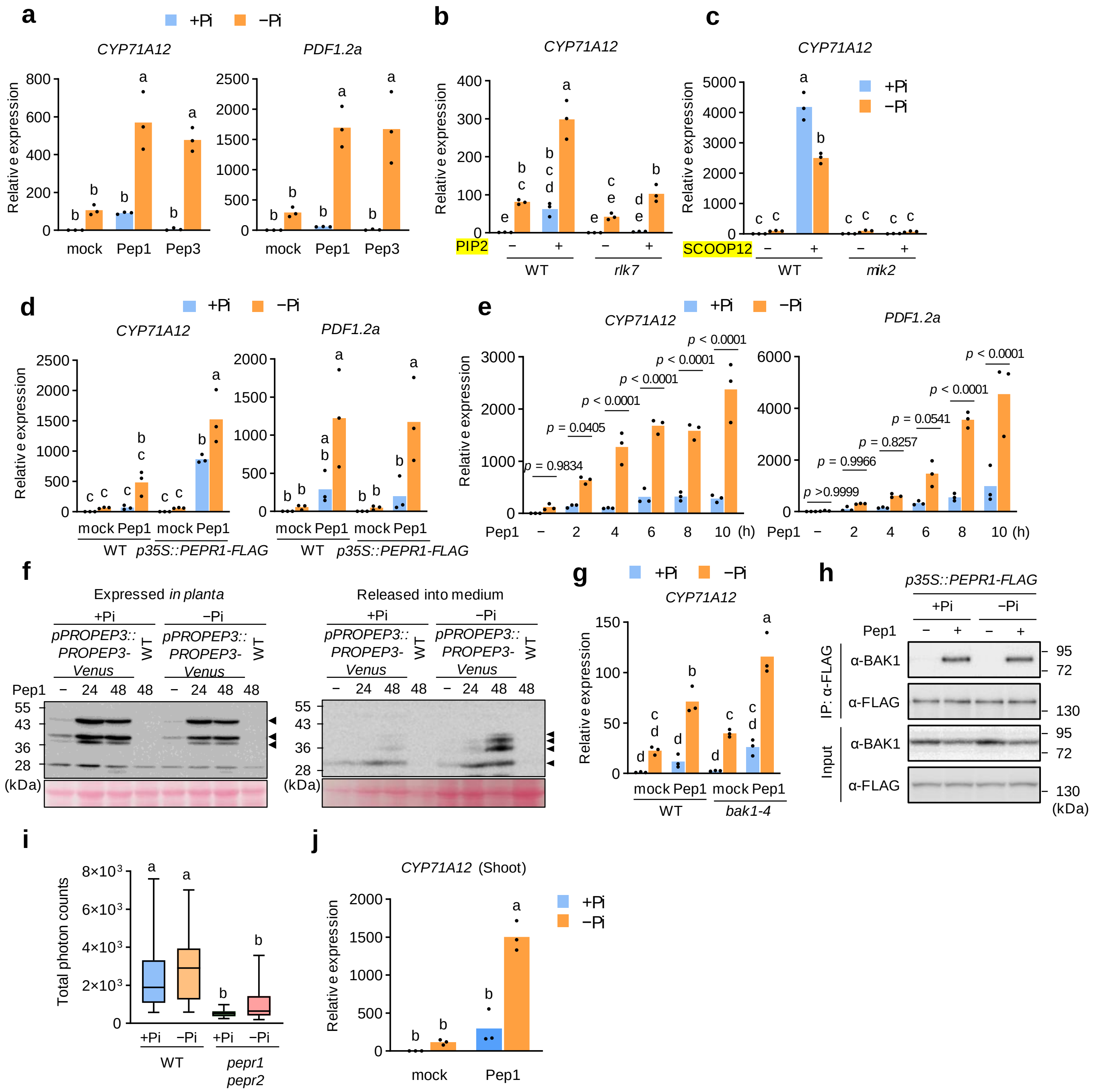


**Supplementary Fig. 3: Pi deficiency enhances multiple PEPR signaling outputs.**
**a**, *CYP71A12* and *PDF1.2a* transcript levels in WT roots grown under +Pi (625 μM) or −Pi (10 μM) and treated with water (mock) or 1 μM Pep3 for 6 h. **b**,**c**, *CYP71A12* transcript levels in WT and *rlk7* (**b**) or *mik2* (**c**) roots grown under +Pi or −Pi and treated with water (mock) or 1 μM PIP2 (**b**) or SCOOP12 (**c**) for 6 h. **d**, *CYP71A12* and *PDF1.2a* transcript levels in WT and *pepr1 pepr2/p35S::PEPR1-FLAG* roots grown under +Pi or −Pi and treated with water (mock) or 1 μM Pep1 for 6 h. **e**, *CYP71A12* and *PDF1.2a* transcript levels in WT roots grown under +Pi or −Pi and treated with 1 μM Pep1 for the indicated times. **f**, Anti-PROPEP3 immunoblot of PROPEP3-Venus in seedlings cultured in liquid medium and treated with or without 1 μM Pep1 for the indicated times. Arrows indicate PROPEP3-Venus; Ponceau S staining is shown as a loading control. **g**, *CYP71A12* and *PDF1.2a* transcript levels in WT and *bak1-4* roots grown under +Pi or −Pi and treated with water (mock) or 1 μM Pep1 for 6 h. **h**, Co-immunoprecipitation of PEPR1-FLAG and BAK1 in transgenic Arabidopsis roots treated with water (mock) or 1 μM Pep1 for 15 min. IP and IB denote

immunoprecipitation and immunoblotting, respectively. **i**, Pep1-induced ROS production in WT and *pepr1 pepr2* seedlings grown under +Pi or −Pi. Box plots show the median, interquartile range, and minimum and maximum (*n* = 20 biological replicates for *pepr1 pepr2* –Pi and *n* = 19 biological replicates for all other treatment groups.). **j**, *CYP71A12* transcript levels in WT shoots grown under +Pi or −Pi and treated with water (mock) or 1 µM Pep1 for 6 h. Different letters indicate significant differences ($p < 0.05$; two-way ANOVA with Sidak's (**a,d**) or Tukey's (**b,c,g,i–j**) multiple comparisons test). In **e**, *p*-values were calculated using two-way ANOVA with Sidak's test).

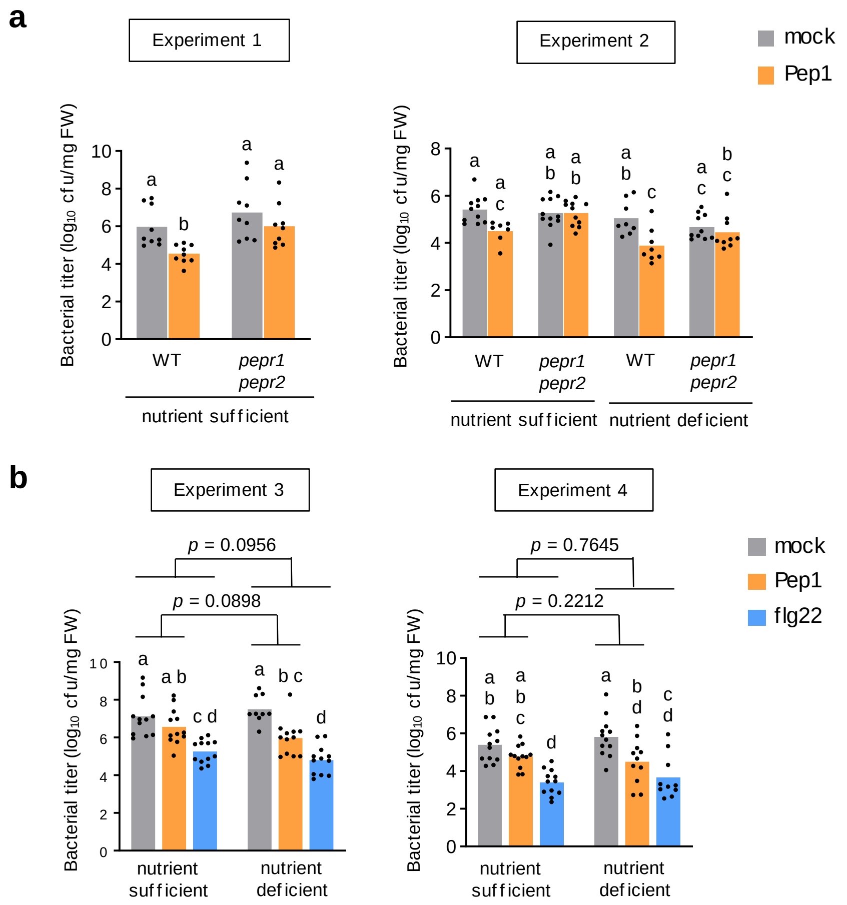


**Supplementary Fig. 4: Pep1- and flg22-induced resistance against *Pto* DC3000 under nutrient sufficient and nutrient deficient conditions.**

**a**,**b**, Growth of *Pto* DC3000 in WT or *pepr1 pepr2* grown on nutrient-sufficient or nutrient-deficient soil. Leaves were infiltrated with water (mock), 1 µM Pep1, or 1 µM flg22 24 h before bacterial inoculation, and bacterial populations were quantified at 3 days post-inoculation. Data are means. In Experiment 1, $n$ = 7, 8, 9, 10, and 11 biological replicates for WT under nutrient-sufficient conditions with Pep1, WT under nutrient-deficient conditions, *pepr1 pepr2* under nutrient-deficient conditions with Pep1, *pepr1 pepr2* under nutrient-deficient conditions with mock, and all other treatment groups, respectively. The data obtained under nutrient-deficient conditions are shown in Fig. 2e. In Experiment 2, $n$ = 11, 8, and 10 biological replicates for mock under nutrient-sufficient conditions, mock under nutrient-deficient conditions, and all other treatment groups, respectively. In Experiment 3, $n$ = 9 and 12 biological replicates for mock under nutrient-deficient conditions and all other treatment groups, respectively. In Experiment 4, $n$ = 10 and 12 biological replicates for flg22 under nutrient-deficient conditions and all other treatment groups, respectively. Different letters indicate significant differences ($p < 0.05$; two-way ANOVA with Tukey's multiple comparisons test). The interaction between nutrient conditions and elicitor treatment was assessed by two-way repeated-measures ANOVA.

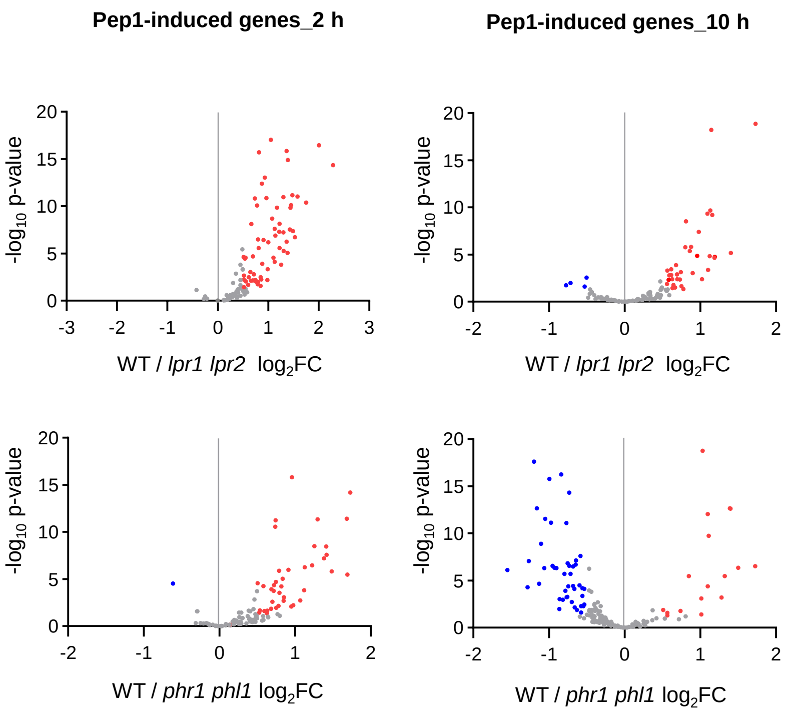


**Supplementary Fig. 5: Expression levels of Pep1-induced genes in *phr1 phl1* and *lpr1 lpr2* mutants.**

Volcano plots showing differentially expressed genes in *lpr1 lpr2* or *phr1 phl1* relative to WT at 2 h and 10 h after Pep1 treatment under low Pi conditions.

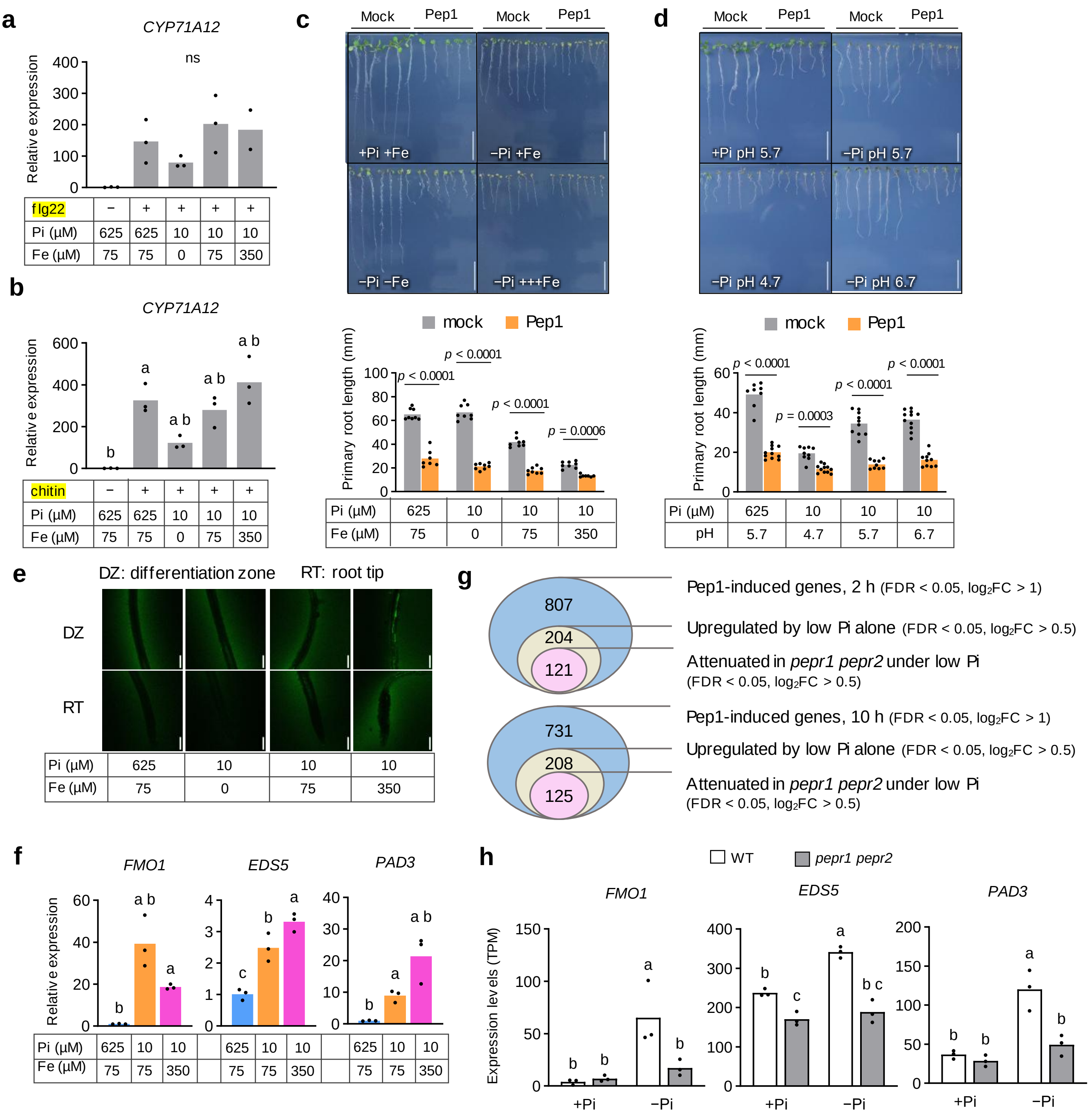


**Supplementary Fig. 6: Pep–PEPR signaling contributes to iron-associated defense activation.** **a,b**, *CYP71A12* transcript levels in WT roots grown under the indicated Pi and Fe conditions and treated with 1 µM flg22 (**a**) or 1 mg ml$^{-1}$ chitin (**b**) for 6 h. Data are means, $n$ = 3 biological replicates. Different letters indicate significant differences ($p$ < 0.05); ns, not significant (one-way ANOVA with Tukey's multiple comparisons test). **c**, Primary root length of Arabidopsis seedlings grown under the indicated Pi and Fe conditions for 7 days with or without 1 µM Pep1. **d**, Primary root length of seedlings grown for 7 days under the indicated Pi and pH conditions with or without 1 µM Pep1. Scale bars, 1 cm. Data are means. In **c**, $n$ = 7, 7, and 8 biological replicates for 625 µM Pi + Pep1, 10 µM Pi + 350 µM Fe + Pep1, and all other conditions, respectively. In **d**, $n$ = 8, 11, 9, 11, 10, 9, 11, and 10 biological replicates for 625 µM Pi (pH 5.7, mock), 10 µM Pi (pH 4.7, Pep1), 10 µM Pi (pH 5.7, mock), 10 µM Pi (pH 6.7, Pep1), 10 µM Pi (pH 4.7, mock), 10 µM Pi (pH 5.7, Pep1), 10 µM Pi (pH 6.7, mock), and 625 µM Pi (pH 5.7, Pep1), respectively. *P*-values were calculated using two-way ANOVA with Sidak's multiple comparisons test). **e**, Carboxy-H2DCFDH staining of ROS in WT seedlings grown under the indicated Pi and Fe conditions. Top, differentiation zone; bottom, root tip. Scale bars, 100 µm. **f**, Transcript levels of defense-related genes in WT roots grown under the indicated Pi and Fe conditions. **g**, Venn diagram of Pep1-induced genes ($log_2FC > 1$, FDR < 0.05) that were upregulated under low Pi ($log_2FC > 0.5$, FDR < 0.05) and downregulated in *pepr1 pepr2* under low Pi. **h**, Transcript levels of defense-related genes in WT and *pepr1 pepr2* grown under +Pi or −Pi. In **f,h**, data are means, $n = 3$ biological replicates; different letters indicate significant differences ($p < 0.05$; two-way ANOVA with Tukey's multiple comparisons test).

**a**

| Group 1 | Group 2 | Sample size | Permutations | pseudo-F | p-value | q-value |
|---|---|---|---|---|---|---|
| WT | *lpr1 lpr2* | 10 | 999 | 3.286156 | 0.009 | 0.0165 |
| WT | *pepr1 pepr2* | 10 | 999 | 2.393201 | 0.013 | 0.0175 |
| WT | *pepr1 pepr2 phr1 phl1* | 9 | 999 | 3.902289 | 0.005 | 0.0165 |
| WT | *phr1 phl1* | 9 | 999 | 3.697142 | 0.009 | 0.0165 |
| WT | *phr1 phl1 lpr1 lpr2* | 9 | 999 | 3.997772 | 0.004 | 0.0165 |
| *lpr1 lpr2* | *pepr1 pepr2* | 10 | 999 | 2.539629 | 0.002 | 0.0165 |
| *lpr1 lpr2* | *pepr1 pepr2 phr1 phl1* | 9 | 999 | 4.117526 | 0.014 | 0.0175 |
| *lpr1 lpr2* | *phr1 phl1* | 9 | 999 | 4.000389 | 0.011 | 0.0165 |
| *lpr1 lpr2* | *phr1 phl1 lpr1 lpr2* | 9 | 999 | 4.583876 | 0.007 | 0.0165 |
| *pepr1 pepr2* | *pepr1 pepr2 phr1 phl1* | 9 | 999 | 3.631585 | 0.011 | 0.0165 |
| *pepr1 pepr2* | *phr1 phl1* | 9 | 999 | 2.392769 | 0.007 | 0.0165 |
| *pepr1 pepr2* | *phr1 phl1 lpr1 lpr2* | 9 | 999 | 4.163685 | 0.009 | 0.0165 |
| *pepr1 pepr2 phr1 phl1* | *phr1 phl1* | 8 | 999 | 1.862718 | 0.032 | 0.032 |
| *pepr1 pepr2 phr1 phl1* | *phr1 phl1 lpr1 lpr2* | 8 | 999 | 2.178542 | 0.03 | 0.032 |
| *phr1 phl1* | *phr1 phl1 lpr1 lpr2* | 8 | 999 | 2.235803 | 0.027 | 0.031154 |

**b**

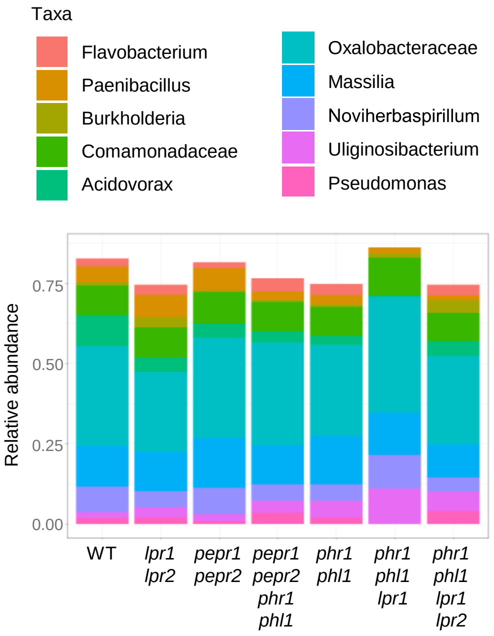


**c** **d**

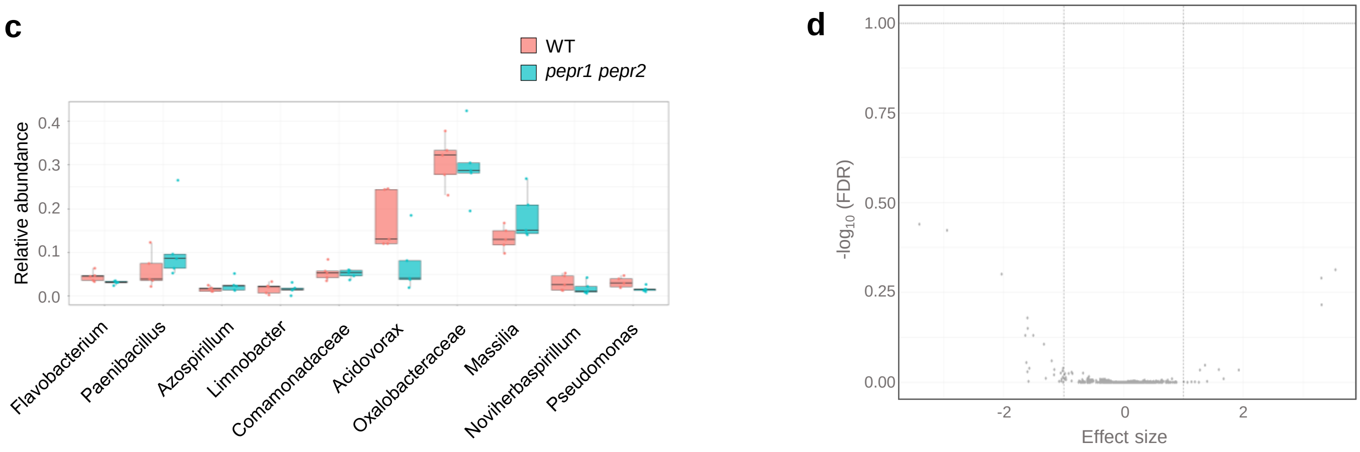


**Supplementary Fig. 7: Community- and taxon-level analyses of the root microbiota under phosphate deficiency.**

**a**, Pairwise PERMANOVA to determine the significance of variables in the root microbiome assembly of different plant genotypes. **b**, Relative abundances of the 10 most abundant bacterial genera in the root microbiota under low Pi conditions. Genera were ranked according to their mean relative abundance across all low-Pi samples. Taxonomic assignments were performed at the genus level using QIIME2.**c**, Relative abundances of the 10 most abundant bacterial genera in the root microbiota of WT and *pepr1 pepr2* grown under low Pi conditions. Box plots show the median, interquartile range, and minimum and maximum ($n$ = 5). **d**, Volcano plot showing differential abundance of amplicon sequence variants (ASVs) between WT and *pepr1 pepr2* plants under low Pi conditions, as determined using ALDEx2. Each point represents one ASV. The x-axis indicates the ALDEx2 effect size, and the y-axis shows the −log10-transformed Benjamini–Hochberg false discovery rate (FDR)-adjusted P value. Horizontal and vertical dashed lines indicate the thresholds used for statistical significance and effect size, respectively.

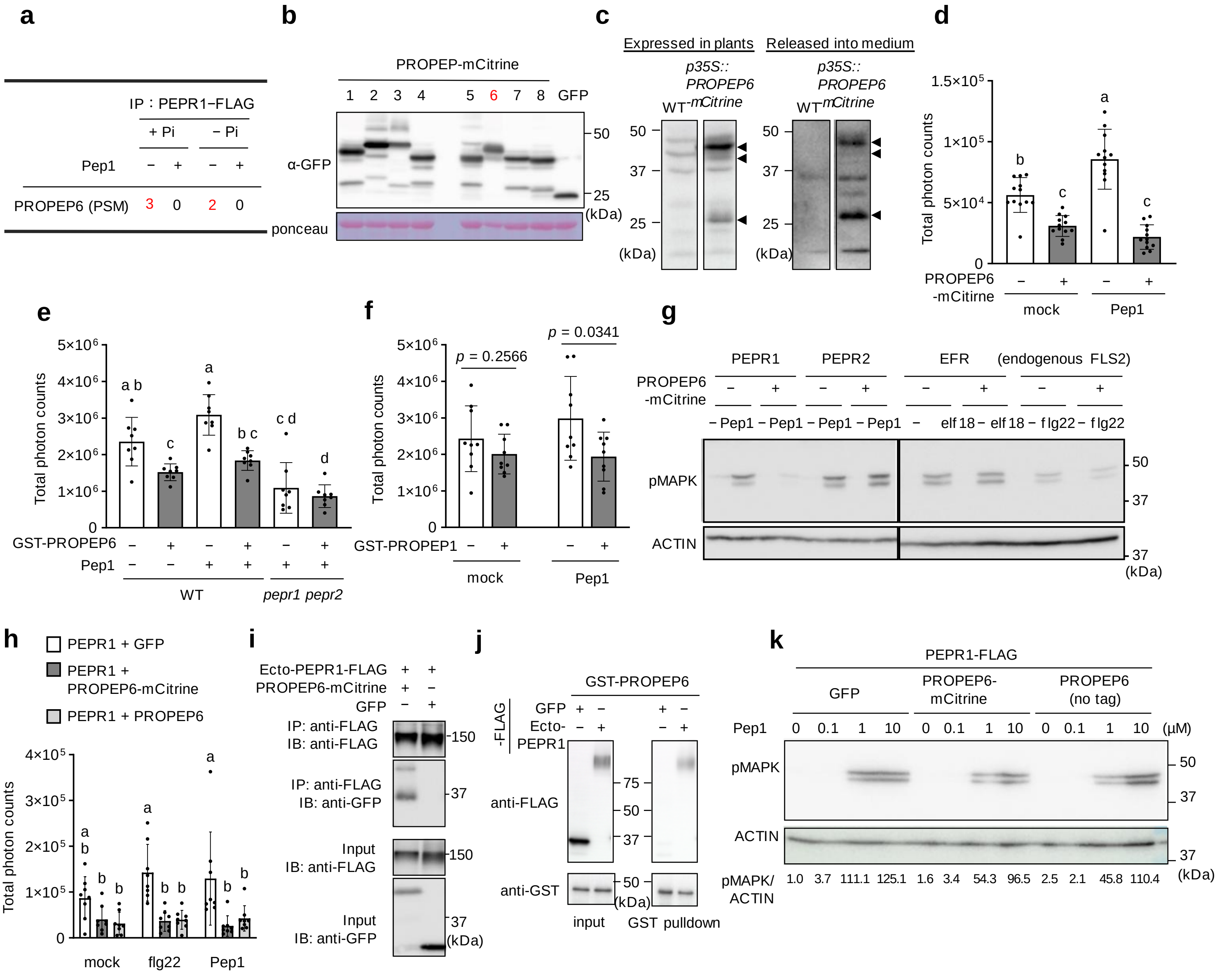


**Supplementary Fig. 8: Characterization of non-canonical PROPEP-derived modulation of PEPR signaling.**

**a**, IP–MS analysis of PEPR1-associated proteins with or without Pep1. Proteins extracted from PEPR1-FLAG transgenic plants were immunoprecipitated with anti-FLAG beads and analyzed by mass spectrometry. Peptide spectrum matches (PSMs) are indicated. **b**, Immunoblot analysis of PROPEP–mCitrine proteins transiently expressed in *N. benthamiana* leaves. **c**, Immunoblot analysis of PROPEP6–mCitrine in *p35S::PROPEP6-mCitrine* seedlings cultured in liquid medium and treated with 1 μM Pep1 for 24 h. Arrows indicate PROPEP6–mCitrine. **d**, ROS production in *N. benthamiana* leaves transiently expressing Arabidopsis PROPEP6. Data are means ± SD, $n$ = 12 biological replicates. **e**, Pep1-induced ROS production in WT and *pepr1 pepr2* leaf discs co-treated with 100 nM Pep1 and 500 nM GST-PROPEP6 or GST. Data are means ± SD, $n$ = 8 biological replicates. In **d**,**e**, different letters indicate significant differences ($p < 0.05$; one-way (**d**) or two-way (**e**) ANOVA with Tukey's multiple comparisons test). **f**, Pep1-induced ROS production in WT leaf discs co-treated with 200 nM Pep1 and 1 μM GST-PROPEP1 or GST. Data are means ± SD, $n$ = 9 biological replicates. *P*-values were calculated using two-sided Student's *t*-test. **g**, MAPK phosphorylation in *N. benthamiana* leaves transiently expressing Arabidopsis PROPEP6 and the indicated receptor after treatment with 1 μM Pep1, elf18, or flg22 for 15 min. ACTIN is served as a loading control. **h**, ROS production in *N. benthamiana* leaves transiently expressing Arabidopsis PROPEP6. Data are means ± SD, $n = 8$ biological replicates. Different letters indicate significant differences ($p < 0.05$; two-way ANOVA with Tukey's multiple comparisons test). **i**, Co-immunoprecipitation of PROPEP6 and Ecto-PEPR1 in *N. benthamiana* leaves expressing the indicated constructs. IP was performed with anti-FLAG beads and PROPEP6 detected with anti-GFP antibody. **j**, Pull-down assay of PROPEP6 and Ecto-PEPR1. Input and output were analyzed by immunoblotting with anti-FLAG and anti-GST antibodies. **k**, MAPK phosphorylation in *N. benthamiana* leaves transiently expressing Arabidopsis PROPEP6 and PEPR1-FLAG after treatment with Pep1 for 15 min. ACTIN is served as a loading control. Numbers below the pMAPK blot indicate relative band intensities normalized to the corresponding ACTIN signals.

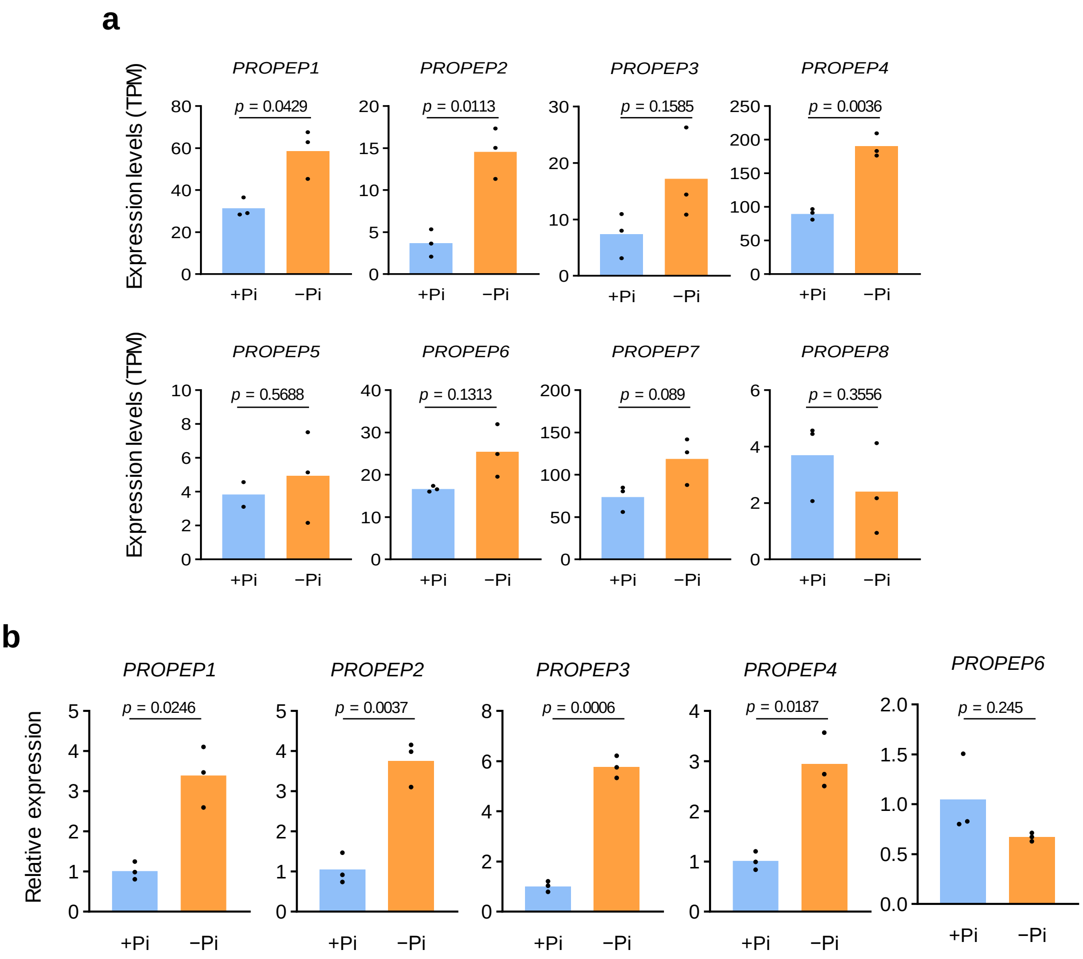


**Supplementary Fig. 9: *PROPEP* expression under different Pi conditions.**

**a**, Transcript levels of *PROPEP* genes in WT roots under Pi-sufficient (+Pi) and Pi-deficient (−Pi) conditions determined by RNA-seq. Data are means, with $n$ = 2 biological replicates for PROPEP5 +Pi, and $n$ = 3 biological replicates for all other conditions. $P$-values were calculated using two-sided Student's $t$-test. **b**, qRT–PCR analysis of *PROPEP* expression in WT roots under +Pi and −Pi conditions. Data are means, $n$ = 3 biological replicates. $P$-values were calculated using two-sided Student's $t$-test).

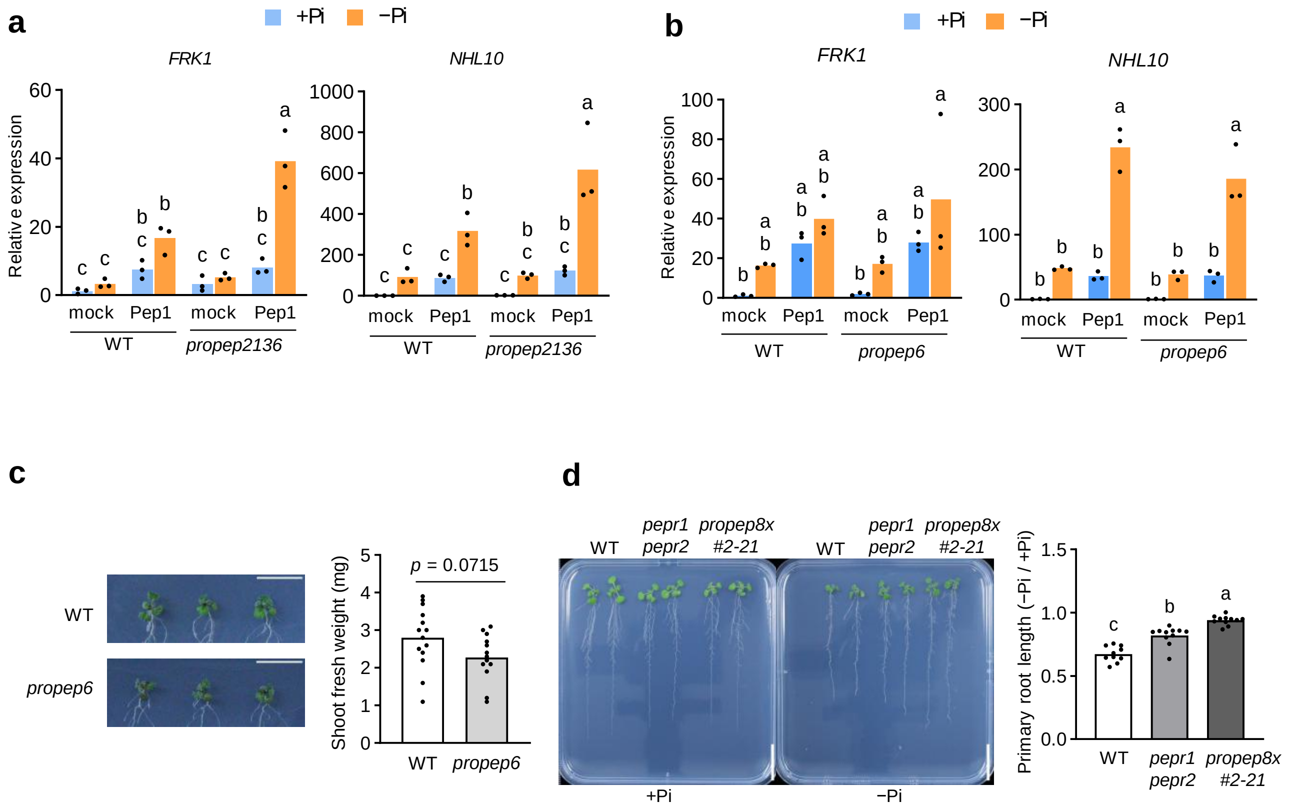


**Supplementary Fig. 10: Genetic analyses of PROPEP-mediated regulation of PEPR signaling and phosphate starvation responses.**

**a**, Transcript levels of defense-related genes in WT and *propep2136* seedlings grown under +Pi or −Pi conditions and treated with 500 nM Pep1 for 6 h. Data are means, $n = 3$ biological replicates. Different letters indicate significant difference ($p < 0.05$; two-way ANOVA followed by Sidak's multiple comparisons test). **b**, Transcript levels of defense-related genes in WT and *propep6* seedlings grown under +Pi or −Pi conditions and treated with 500 nM Pep1 for 6 h. Data are means, $n = 3$ biological replicates. Different letters indicate significant difference ($p < 0.05$; two-way ANOVA followed by Sidak's multiple comparisons test). **c**, Three-day-old seedlings were exposed to Pi-deficient conditions (10 µM Pi) for 10 days. Scale bar, 20 mm. Data are means, $n = 14$ biological replicates for WT and $n = 13$ biological replicates for *propep6* biological replicates. *P*-values were calculated using two-sided Student's *t*-test. **d**, Primary root length of Arabidopsis seedlings grown under +Pi (625 µM) or −Pi (50 µM) conditions. Scale bar, 20 mm. Data are means, $n = 10$ biological replicates for WT and $n = 11$ biological replicates for all other mutants. Different letters indicate significant differences ($p < 0.05$; two-way ANOVA followed by Tukey's multiple-comparisons test).
